# Sensitization of FOLFOX-resistant colorectal cancer cells via the modulation of a novel pathway involving protein phosphatase 2A

**DOI:** 10.1101/2021.07.21.453259

**Authors:** Satya Narayan, Asif Raza, Iqbal Mahmud, Nayeong Koo, Timothy J. Garrett, Mary E. Law, Brian K. Law, Arun K. Sharma

## Abstract

The treatment of colorectal cancer (CRC) with FOLFOX shows some efficacy, but these tumors quickly develop resistance to this treatment. We have observed an increased phosphorylation of AKT1/mTOR/4EBP1 and levels of p21 in FOLFOX-resistant CRC cells. We have identified a small molecule, NSC49L, that stimulates protein phosphatase 2A (PP2A) activity, downregulates the AKT1/mTOR/4EBP1-axis, and inhibits p21 translation. We have provided evidence that NSC49L- and TRAIL-mediated sensitization is synergistically induced in p21-knockdown CRC cells, which is reversed in p21-overexpressing cells. p21 binds with procaspase 3 and prevents activation of caspase 3. We have shown that TRAIL induces apoptosis through the activation of caspase 3 by NSC49L-mediated downregulation of p21 translation, and thereby cleavage of procaspase 3 into caspase 3. NSC49L does not affect global protein synthesis. These studies provide a mechanistic understanding of NSC49L as a PP2A agonist, and how its combination with TRAIL sensitizes FOLFOX-resistant CRC cells.

**Graphical Abstract:** 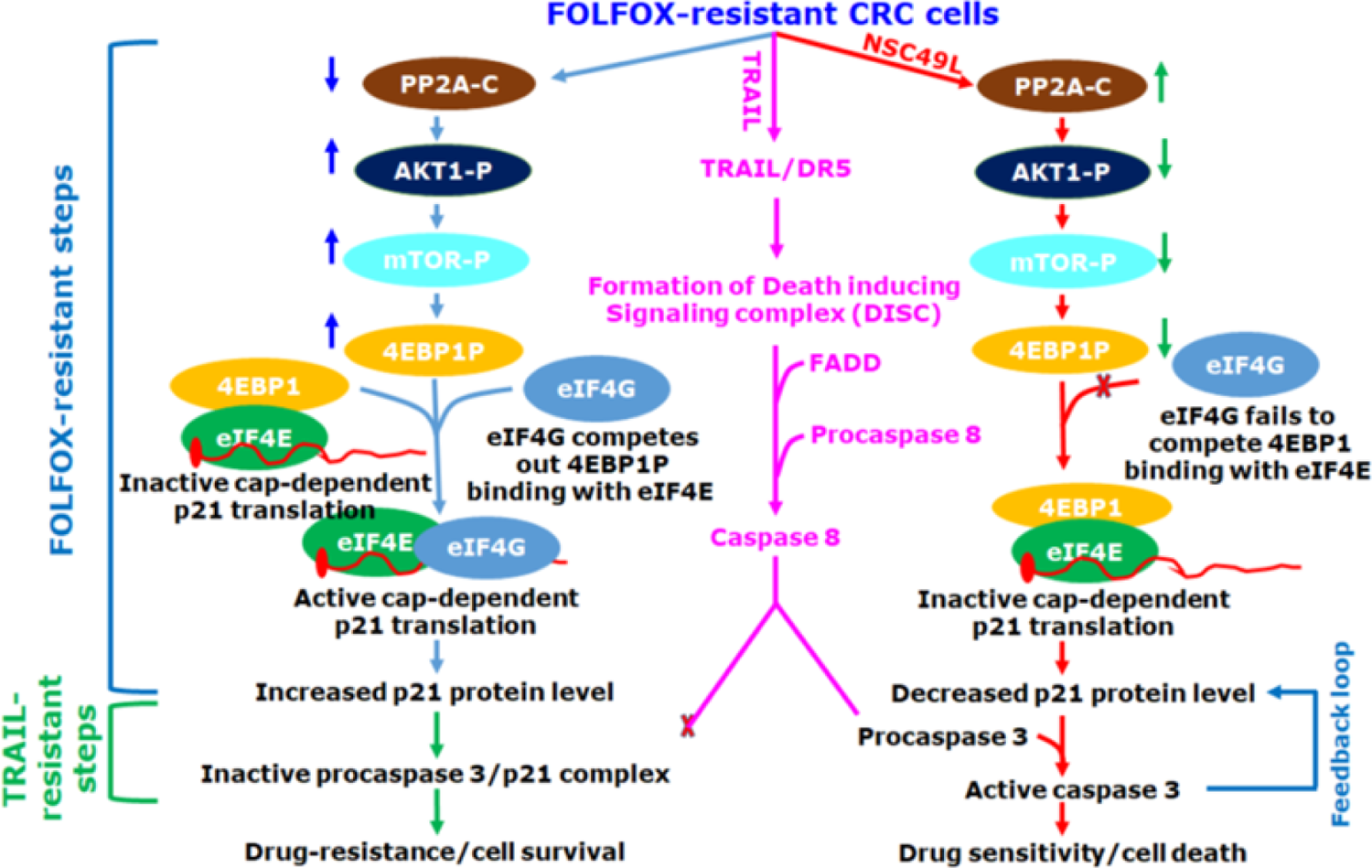

## INTRODUCTION

Colorectal cancer (CRC) is the third most common cancer and the second leading cause of cancer death for men and women in the USA. The American Cancer Society has estimated about 147,950 new cases of CRC with an estimated 53,200 deaths in both sexes in the year 2020 (Siegel et al., 2020). Modern CRC therapy has achieved considerable progress with the use of aggressive surgical resection and chemotherapy. However, nearly 50% of patients with colonic tumors develop recurrent metastatic disease (Young et al., 2014). 5-Fluorouracil (5-FU) is the most integral systemic component of curative and palliative therapy for CRC, but the overall response rate in advanced CRC is only 10-15%. The current clinical practice utilizes the combination of 5-FU with folinic acid and oxaliplatin (FOLFOX) as a first-line chemotherapy for metastatic CRC; however, CRC tumors quickly develop resistance to FOLFOX, which carries a significant toxicity, cost, and patient morbidity (Yaffee et al., 2015). Despite the use of chemotherapy (oxaliplatin-based: FOLFOX or CAPOX, irinotecan-based: FOLFIRI) combined with Bevacizumab (anti-VEGF), Cetuximab (anti-EGFR), or panitumumab (for RAS wild-type tumors) to manage metastatic CRC tumors, the five-year survival rate remains low at 14% (Ikoma et al., 2017), which emphasizes the urgent need for a better second-line chemotherapy for advanced stage CRC.

At a molecular level, the activation of AKT1 signaling has been reported in 57% of CRCs as an early event during sporadic colon carcinogenesis, which can occur through gene amplification, mutation or epigenetic manipulation (Roy et al., 2002). The increased levels of phosphorylated AKT1, mammalian target of rapamycin (mTOR), and eukaryotic translation initiation factor 4E-binding protein 1 (4EBP1) are linked with drug-resistance and poor prognosis of CRC patients (Li, 2018; Malinowsky et al., 2014; Melling et al., 2015). In the past few years, the development of several AKT1/mTOR/4EBP1-axis inhibitors have been described, but the success rate has been unsatisfactory (Xu et al., 2016). The discovery of perifosine (AKT1 inhibitor) in combination with capecitabine is showing some promise with CRC therapy (Richardson et al., 2012). The mTOR inhibitors have not been very successful for CRC therapy due to the activation of upstream PI3K activity (Rozengurt et al., 2014). MAPK (Hoang et al., 2012) and EGFR (Wei et al., 2015) activation and 4EBP1 phosphorylation (T37/46P) prevention (Gerdes et al., 2013) are also reported with AKT1 and mTOR inhibitors. Further, activation of the AKT1 pathway also enhances the function of eukaryotic translation initiation factor 4E (eIF4E), which selectively increases the translation of key mRNAs involved in tumor growth, angiogenesis, and cell survival (Graff et al., 2008). Besides the mentioned problems above, these therapeutic compounds are toxic to normal cells.

A possible mechanism to decrease drug-resistance is via the dephosphorylation of AKT1 by PP2A, a tumor suppressor, belonging to the family of protein serine/threonine phosphatases (Ruvolo, 2016). PP2A is a heterotrimeric holoenzyme with three subunits: PP2A-C (catalytic), PP2A-B (regulatory) and PP2A-A (scaffolding). Each subunit further exists in multiple isoforms, such as PP2A-C exists as two isoforms – PP2A-CA and PP2A-CB (Janssens et al., 2008).

Frequent inactivation of PP2A in CRC cell lines and patient tumor samples has been documented (Cristobal et al., 2014; Wang et al., 1998). In a recent clinical study, low expression of PP2A was linked with worse overall survival for CRC patients, and was suggested as an independent prognostic factor for CRC (Yong et al., 2018). In recent years, there has been a great focus towards the discovery of PP2A agonists for the downregulation of oncogenic phosphorylation of kinases, implicating PP2A as a useful target for therapeutic development (Mazhar et al., 2019; O’Connor et al., 2018; Sangodkar et al., 2017; Tohme et al., 2019; Westermarck, 2018). While a significant success has been achieved with new PP2A activators targeting to different tumor types, studies with the FOLFOX-resistant CRC tumors are not yet reported. Further, the *in vivo* toxicity of these PP2A activators remains a concern.

Tumor necrosis factor (TNF)-related apoptosis-inducing ligand (TRAIL) is a member of a subset of the TNF receptor superfamily of protein ligands, which are a physiological means of killing many cancer cells, including CRC cells, while sparing the normal cells (Ashkenazi et al., 1999). Thus, TRAIL has great promise as a tumor-selective anti-cancer agent. Unfortunately, the major limitation of the TRAIL therapy is the development of resistance in cancer cells, comprising CRC cells (Van der Jeught et al., 2018). TRAIL, a trimeric protein, binds with death receptor (DR) molecules and forms an assembly platform for the adaptor protein Fas-associated death domain (FADD) (Ralff and El-Deiry, 2018). Subsequently, FADD binds to procaspase 8, and together with DR and procaspase 8 forms the death-inducing signaling complex (DISC) that promotes activation of caspase 8. The activated caspase 8 is released into the cytosol, where it activates the executioner caspase 3 that drives apoptotic breakdown of the cell (Carneiro and El-Deiry, 2020). Hence, the inactivation of one of these steps can contribute to TRAIL- or drug-resistance via evasion of apoptosis. In previous studies, the interaction of p21 with procaspase 3 has been shown to block the cleavage of procaspase 3 into executioner caspase 3 that may prevent the executioner caspase 3 from inducing apoptosis (Suzuki et al., 2000). Increased levels of p21 have been identified as a drug-resistance factor in several cancer cells (Georgakilas et al., 2017), including 5-FU resistance to CRC cells (Maiuthed et al., 2018).

In the present study, we have identified p21 as an acquired FOLFOX resistance factor in CRC cells. We report that the increased FOLFOX-resistant CRC cells have increased levels of p21, which is independent of p53-mediated transcription. We have discussed the role of a novel PP2A agonist, NSC49L [NSC30049 ((1-(4-Chloro-2-butenyl)-1λ∼5∼,3,5,7-tetraazatricyclo [3.3.1.1∼3,7∼]decane)] that by inhibiting the AKT1/mTOR/4EBP1 pathway, reduces p21 translation. The decreased level of p21 released procaspase 3 for TRAIL-dependent synergistic sensitization of FOLFOX-resistant CRC cells. Thus, downregulation of p21 can be a critical mechanism for abolishing drug-resistance and inducing TRAIL-mediated apoptosis in FOLFOX-resistant CRC cells.

## RESULTS

### *PPP2CB* was augmented in CRC tumors and closely associated with poor prognosis in CRC patients

We used the Cancer Genome Atlas (TCGA) through the Genomic Data Commons Data Portal for determining the expression level of *PPP2CB* (catalytic subunit B) in CRC tumors compared to matched normal tissues. *PPP2CB* was significantly and steadily decreased during the progression from stage I to stage IV in CRC tumors compared to adjacent normal tissues (Fig. 1A). We also analyzed *PPP2CB* levels in CRC cells spread in lymph nodes. There was a progressive significant decrease in the *PPP2CB* levels at different nodal stages (N0 to N2) of CRC development compared to normal tissues (Fig. 1A). A similar decreased pattern of *PPP2CB* expression was also identified in mucinous and non-mucinous CRC tumors compared to normal tissues (Fig. S1A). Then, we used the Clinical Proteomic Tumor Analysis Consortium (CPTAC) data portal and found a decreased PP2AC protein level in CRC tumors compared to normal tissues (Fig. 1B, S1B). Further, the low-level expression of *PPP2CB* was associated with poor prognosis of CRC patients (Fig. 1C, S1C). These results suggest a negative correlation of *PPP2CB* expression with CRC progression and poor prognosis of CRC patients.

**Figure 1.**
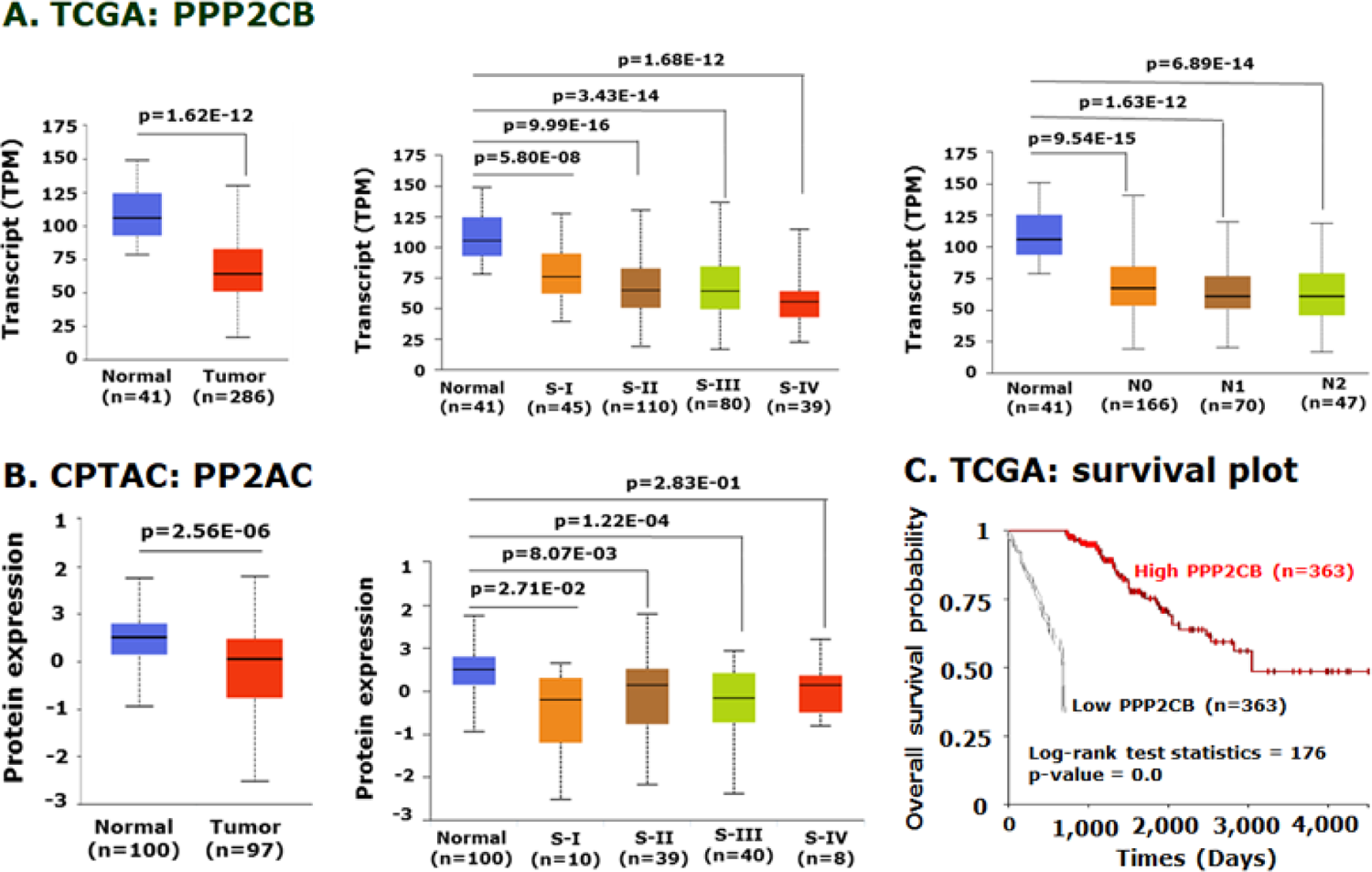
PP2AC expression level is directly correlated with disease progression and the survival of CRC patients. **A**, Box plot depicting a summary of TCGA data quantification of PPP2CB in normal tissue and CRC tumors (left), different stages (middle), and different nodal progression (right). **B,** Box plot showing the CPTAC data quantitation of PP2AC protein levels in normal tissues and CRC tumors (left), and different stages of progression (right). **C,** Kaplan-Meier survival analysis of TCGA data was plotted for the CRC cases that were high and low in PPP2CB expression. The p-value is shown on the top of each graph.

### *PPP2CB* expression is negatively linked with the expression of *AKT1*, *mTOR*, *4EBP1* and *CDKN1A* in FOLFOX-responsive and non-responsive CRC tumors

Since AKT1 is a downstream target of PP2A and mTOR, 4EBP1 and p21 are on the same axis of regulation, we determined the correlation of their gene expression in FOLFOX-responsive and non-responsive CRC tumors. In this study, all publicly available databases were searched and three gene expression datasets of CRC patients with FOLFOX therapy were combined, as described in an earlier study (Lin et al., 2018). We found that the *PPP2CB* gene expression level was significantly higher in FOLFOX-responsive versus non-responsive tumors (Fig. 2). On the other hand, the gene expression level of *AKT1*, *mTOR*, *4EBP1* and *CDKN1A* were significantly lower in FOLFOX-responsive versus non-responsive tumors (Fig. 2), suggesting a negative correlation of *PPP2CB* with *AKT1*, *mTOR*, *4EBP1*and *CDKN1A* in FOLFOX-responsive versus non-responsive CRC tumors. Since increased expression of *AKT1* (Surov et al., 2018), *mTOR* (Wu et al., 2018) and *4EBP1* (Malinowsky et al., 2014) are linked with poor prognosis of CRC patients, and linked to increased levels of p21 with increased proliferative capacity of colon cancer cases (Palaiologos et al., 2019), we hypothesized that the upregulation of the PP2A activity and the downregulation of the AKT1, mTOR, 4EBP1 and p21 levels in CRC cells can be a suitable means of inducing the sensitization of the CRC tumors.

**Figure 2.**
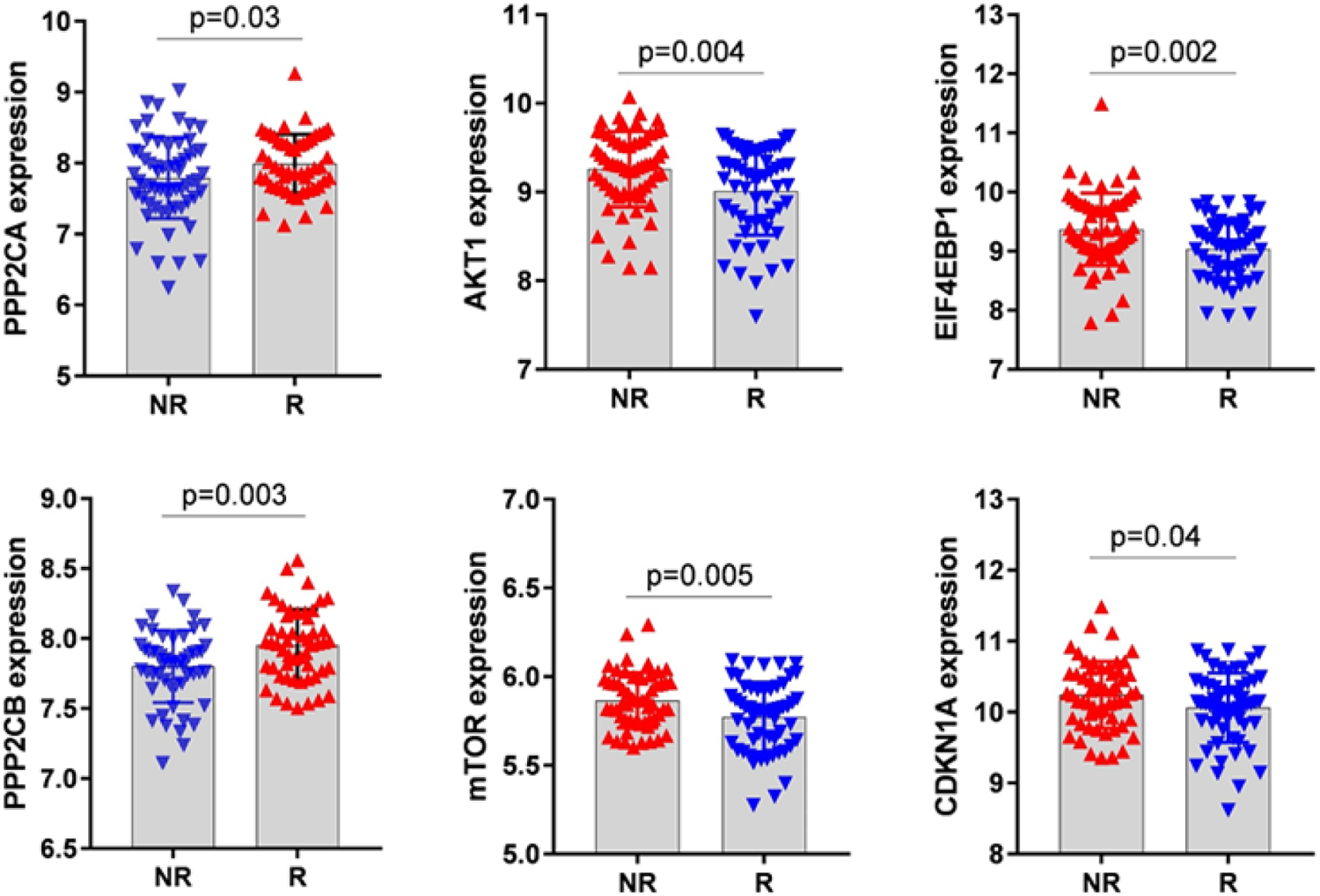
*PPP2CA/B* expression is lower in FOLFOX-non-responsive (NR) versus responsive (R) CRC tumors, which negatively correlated with the expression of *AKT1*, *mTOR*, *4EBP1* and *CDKN1A*. Data was analyzed from the NCBI GEO accession number GSE72970. Number of samples in the NR and R groups were 61 and 63, respectively. The p-value is shown on the top of each graph.

### NSC49L activates PP2A

We have been investigating a small molecule, NSC49L, to understand the molecular mechanisms responsible for its anti-cancer activity. We examined whether it is a kinase inhibitor but the results of a panel of 366 kinases showed that it was not a kinase inhibitor (Narayan et al., 2019). However, from these experiments it appeared that NSC49L might be a phosphatase agonist, as it inhibited hydroxyurea- and 5-FU-induced phosphorylation of Chk1(S317P) and Chk1(S296) in HCT116 and HT29 cell lines (Narayan et al., 2017; Narayan et al., 2019). In previous studies, the involvement of PP2A has been suggested in the ATR-dependent dephosphorylation of Chk1 at S317 and S296 residues (Leung-Pineda et al., 2006).

Another link between NSC49L and Chk1 may be through the downregulation of mTORC1 activity by the downregulation of AKT1 activity, and in turn, the downregulation of the Chk1 phosphorylation (Zhou et al., 2017). The later possibility prompted us to examine whether NSC49L is a PP2A agonist that downregulates the AKT1/mTOR/4EBP1-axis in FOLFOX-resistant CRC cells.

First, we performed an *in vitro* PP2A activity assay using recombinant PP2A-Cα protein and threonine phosphopeptide (KRpTIRR) as substrate in a Malachite green assay (Lek et al., 2017). NSC49L stimulated PP2A-Cα activity in a dose-dependent manner with a Kact(NSC49L) of 14 nM and Vmax of 3384 nmoles/mg/min (Fig. 3A and B). We also determined the effect of NSC49L treatment on the stimulation of PP2A-Cα in FOLFOX-HT29 cells. The immunocomplexes from the control and treated groups were isolated with anti-PP2A-C antibody and were used for PP2A-Cα activity (Lek et al., 2017). Results showed an increased PP2A activity in a dose-dependent manner (Fig. 3C), suggesting that NSC49L acts as a PP2A agonist, which stimulates its activity at nanomolar concentrations.

**Figure 3.**
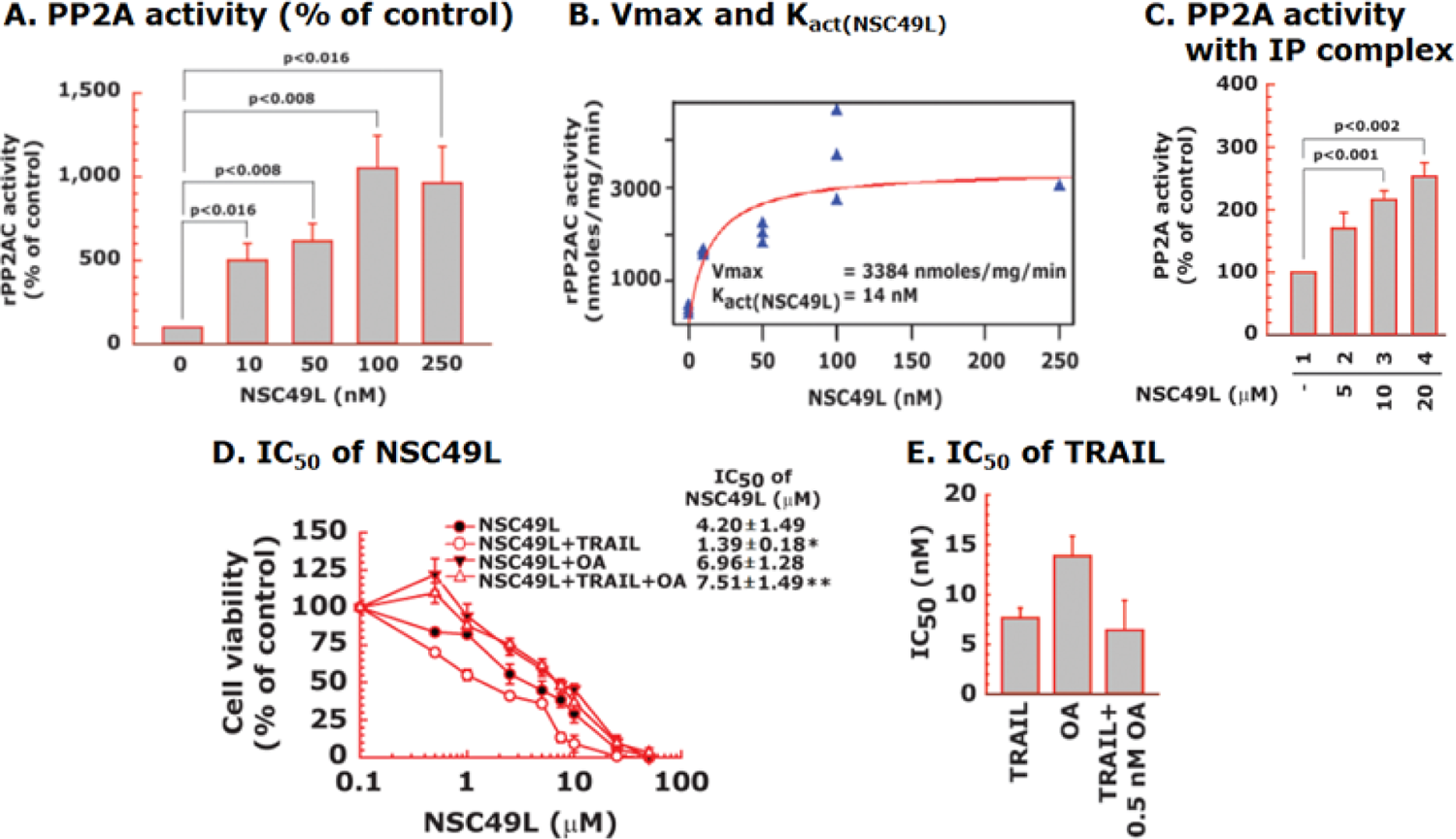
NSC49L-mediated activation of PP2AC. **A**, human rPP2AC activity determined with the malachite green assay. **B,** Kact(NSC49L) and Vmax values of NSC49L with PP2AC. **C,** PP2A activity using immunecomplexes isolated with anti-PP2AC antibody from control and NSC49L-treated FOLFOX-HT29 cells. Data are the mean ± SE of three experiments. **D,** Okadaic acid (OA), a PP2A inhibitor, blocks NSC49L and TRAIL-induced cytotoxicity to FOLFOX-HT29 cells. Cells were treated with different concentrations of NSC49L, TRAIL and OKA either alone are in combination as indicated in the figure. After 72 h treatment, cell viability was determined using the MTT-assay. **E,** effect of OA on the IC50 of the TRAIL. Data are the mean ± SE of four estimations. The p<0.05 is considered significant.

Okadaic acid (OA) is a PP2A and PP1 inhibitor, but at a selective concentration of 2 nM or lower, it inhibits only PP2A activity (Lu et al., 2009). To determine whether the inhibition of PP2A activity by OA increases cell viability, we treated FOLFOX-HT29 cells with a fixed concentration of OA (0.5 nM) and TRAIL (1 nM) and with different concentrations of NSC49L. Then, we determined the IC50 of NSC49L by MTT-cell viability assays. Results showed that treatment with 0.5 nM of OA either alone or in combination with 1 nM of TRAIL increased the survival of FOLFOX-HT29 cells, as is evident from the increased IC50 of NSC49L (Fig. 3D).

The concentration of OA (0.5 nM) used in this experiment was much lower than its IC50 (∼15 nM). OA at the lower concentration did not affect the IC50 of the TRAIL (Fig. 3E). These results suggest that NSC49L mediates its effect through the stimulation of the PP2A activity in FOLFOX-HT29 cells, which is further effective when treated in combination with TRAIL.

### NSC49L and TRAIL treatments sensitize FOLFOX-resistant HCT116 and HT29 cell lines

HCT116 and HT29 are TRAIL-sensitive and TRAIL-resistant CRC cell lines, respectively (Kim et al., 2015). NSC49L shows more toxicity to HCT116 cells than to HT29 cells, which were further sensitized after TRAIL treatment. The normal colonic epithelial cell line, FHC, was resistant to TRAIL (Fig. 4A). To examine the effect of TRAIL on FOLFOX-resistant CRC cells, we used isogenic FOLFOX-resistant cells derived from HCT116 and HT29 cell lines (Narayan et al., 2017; Yu et al., 2009). TRAIL reduced the viability of both FOLFOX-HCT116 and FOLFOX-HT29 cell lines, while the TRAIL-resistant HT29-derived FOLFOX-HT29 cells were more sensitive than the TRAIL-sensitive HCT116-derived FOLFOX-HCT116 cells (Fig. 4A).

**Figure 4.**
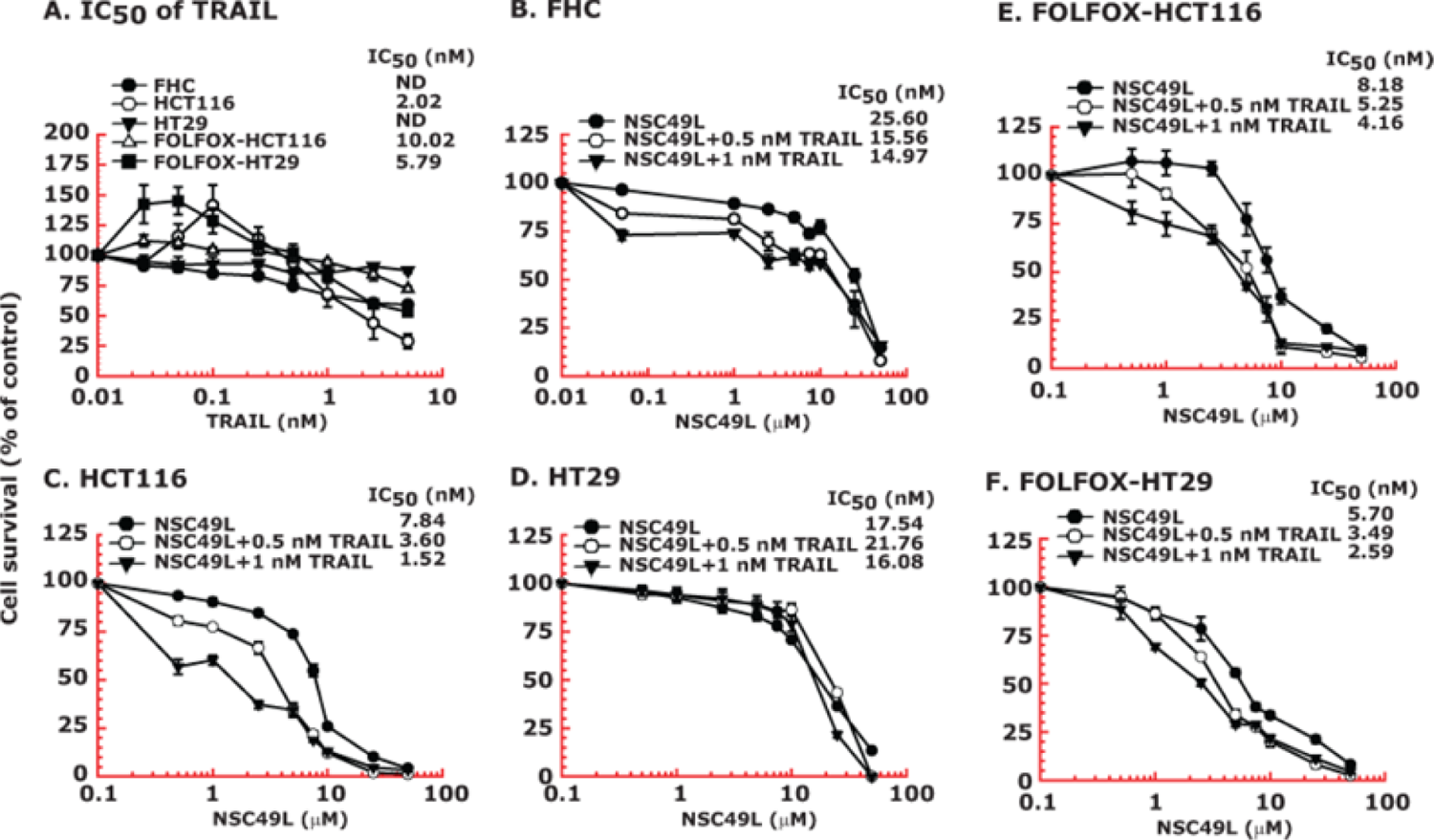
FOLFOX-HT29 and FOLFOX-HCT116 cell lines are much more sensitive than normal colonic epithelial cell line (FHC) to NSC49L and TRAIL treatment. Cells were grown in 96-well plates and treated for 72 h with different concentrations of NSC49L and TRAIL either alone or in combination. Survival of cells was determined by the MTT-assay. **A,** IC50 of TRAIL. **B-F,** IC50 of NSC49L either alone or in the presence of TRAIL in different CRC cell lines. Data are the mean ± SE of four estimations.

Then, we determined whether TRAIL in combination with NSC49L can enhance the cytotoxicity of FHC, HCT116, HT29, FOLFOX-HCT116 and FOLFOX-HT29 cell lines. The results showed little toxicity of NSC49L to FHC and HT29 cells below 10 µM, which was increased in combination with TRAIL, but to a much lower extent than to HCT116 cells (Fig. 4B, C and D). On the other hand, NSC49L sensitized both FOLFOX-HCT116 and FOLFOX-HT29 cell lines to TRAIL (Fig. 4E and F). However, treatment with NSC49L in combination with TRAIL enhanced the FOLFOX sensitivity of FOLFOX-HCT116 and FOLFOX-HT29 cell lines. We observed a dose-dependent decrease in the IC50 (an indicator of increased sensitization or cytotoxicity) of NSC49L in the presence of TRAIL in both FOLFOX-resistant cell lines (Fig. 4E and F).

### NSC49L and TRAIL treatments downregulate AKT1(S473P) and p21 levels in FOLFOX-HCT116 and FOLFOX-HT29 cell lines

Once we established that NSC49L stimulates PP2A activity, we examined its effect on PP2A expression levels. We performed RNAseq analysis of control and NSC49L/TRAIL-treated FOLFOX-HT29 cells. The results showed an increased expression of *PPP2CA* and *PPP2CB*, which are the catalytic subunits of PP2A (Fig. 5A).

**Figure 5.**
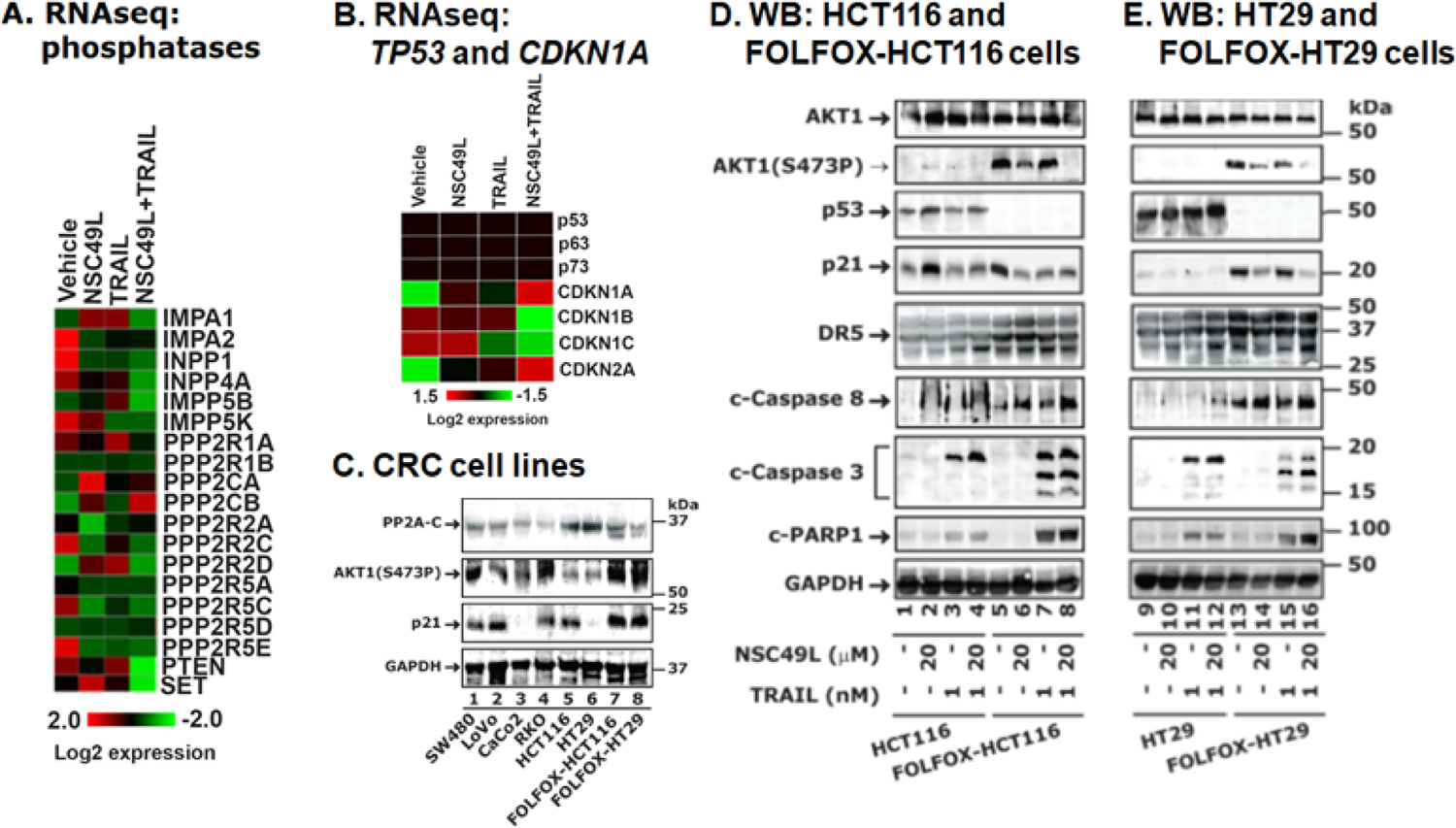
RNAseq and protein analysis of PP2A and downstream target proteins in CRC cells. **A** and **B,** RNAseq analysis of FOLFOX-HT29 cells. Cells were treated with 20 µM of NSC49L and 1 nM of TRAIL either alone or in combination for 16 h. Total RNA was isolated by Trizole reagent and processed for RNAseq analysis. Ward clustering algorithm with euclidean distance measurement were used for heat map analysis. **C,** correlation of PP2A expression level with AKT1(S473P) and p21 levels in CRC cell lines. To determine a correlation of PP2A with AKT1(S473P) and p21, we performed western bots of these protein using the whole cell lysate of different CRC cell lines, including FOLFOX-resistant HCT116 and HT29 cells. **D and E,** protein levels of AKT1, p53, p21, DR5, caspase 3, caspase 8 and PARP1 in HCT116, FOLFOX-CHT116, HT29 and FOLFOX-HT29 cell lines, respectively, treated with NSC49L and TRAIL either alone or in combination for 24 h. The blots are representative of three experiments.

However, the expression of other phosphatases was rather decreased as compared to treated groups. We also performed RNAseq analysis of *TP53* and *CDKN1A* and other related genes. We found that the expression of *TP53*, *TP63* and *TP73* (also known as *p53*, *p63* and *p73*, respectively) was absent in untreated FOLFOX-HT29 cells which was unchanged after the NSC49L or TRAIL treatments (Fig. 5B). Whether the decreased expression level of *p53*, *p63* and *p73* in these cells after NSC49L/TRAIL treatment was regulated at the level of transcription or mRNA degradation is currently unknown. Further, the expression of *CDKN1A* (*p21Cip1/Kip1*) and *CDKN2A* (*p16INK4a*) was increased but *CDKN1B* (*p27Kip1*) and *CDKN1C* (*p57Kip2*) was decreased (Fig. 5B). While the expression of p21 and p16 are linked with drug-resistance (Kreis et al., 2019), we focused on p21 due to its involvement in 5-FU resistance in CRC cells (Maiuthed et al., 2018).

We determined if there is any correlation between PP2AC, AKT1(S473P) and p21 levels in different CRC cell lines that are either resistant or sensitive to FOLFOX. The results showed a negative correlation between the PP2AC/AKT1(S473P) and PP2AC/p21 protein levels in most of the CRC cell lines, but a significant correlation was evident in FOLFOX-HCT116 and FOLFOX-HT29 cell lines (Fig. 5C and S2). These results suggest that, at least, in FOLFOX-HCT116 and FOLFOX-HT29 cell lines, the reduced levels of PP2AC are clearly correlated with increased levels of AKT1(S473P) and p21.

Next, we determined whether the NSC49L and TRAIL-mediated sensitization of FOLFOX-HCT116 and FOLFOX-HT29 cell lines, as observed in Fig. 4, are linked with increased PP2A activity and decreased AKT1(S473P) and p21 levels. We simultaneously compared these results with HCT116 and HT29 cell lines to compare the effects between the FOLFOX-HCT116 and FOLFOX-HT29 cell lines. In untreated HCT116 and HT29 cell lines we did not observe any increase in the level of AKT1(S473P) (Fig. 5D and E, compare lanes 1-4, and 9-12, respectively); however, it was highly increased in FOLFOX-HCT116 and FOLFOX-HT29 cell lines, which was decreased after NSC49L and TRAIL treatments (Fig. 5D and E, compare lanes 5-8, and 13-16, respectively). The expression of p53 was seen in both HCT116 and HT29 cell lines, which was completely absent in FOLFOX-HCT116 and FOLFOX-HT29 cell lines (Fig. 5D and E). Whether the silencing of p53 in these cell lines is at the transcription or translation level is currently not known. Since p21 is regulated by p53, we observed an increase in p21 in HCT116 cells that carry a wild-type *TP53* gene, but not in HT29 cells that carry a mutant *TP53* gene. Surprisingly, we observed a p53-independent increase in the level of p21 in FOLFOX-HCT116 and FOLFOX-HT29 cell lines, which was decreased after NSC49L and TRAIL treatments (Fig. 5D and E, compare lanes 5-8 and 13-16, respectively). Thus, these results suggest that AKT(S473P) and p21 are associated with resistance of FOLFOX-HCT116 and FOLFOX-HT29 cell lines after 5-FU and oxaliplatin treatments. The level of cleaved caspase 8, caspase 3 and PARP1, indicators of apoptosis, were also more prominent when NSC49L and TRAIL were combined (Fig. 5D and E, compare lanes 5-8 and 13-16, respectively). These results suggest that NSC49L-induced PP2A activity and the TRAIL-induced apoptotic pathway are linked with increased caspase 8, caspase 3, and PARP1 levels in FOLFOX-HCT116 and FOLFOX-HT29 cells. These results are supported by a TCGA analysis in which we found a positive correlation between PPP2CB expression and caspase8, caspase 3, and PARP1 expression (Fig. S3).

Since the activation of PP2A by NSC49L dephosphorylates AKT1, the effect of TRAIL on AKT1 dephosphorylation of these cells could be indirect. This observation is similar to TRAIL-induced sensitization of prostate cancer cells in response to the PIP3 phosphatase PTEN, suggesting that activation of the AKT1 pathway may also be a common event in TRAIL resistant cells (Xu et al., 2010). How TRAIL might induce its effect through the AKT1 pathway is not clearly understood. In these studies, we observed an increased effect of NSC49L on AKT1 dephosphorylation when combined with TRAIL. Hence, we expected that the increased expression of DR5, a TRAIL receptor, in FOLFOX-HCT116 and FOLFOX-HT29 cell lines might have been one of the reasons for TRAIL/NSC49L-induced sensitization of these cells (Fig. 5D and E, compare lanes 1 with 5 and 9 with 13, respectively). To understand if the low level of DR5 in HT29 cells might have been the reason for non-responsiveness to TRAIL (Fig. 4A), we overexpressed tetracycline (tet)-inducible DR5 (Fig. 6A) in these cells and determined the sensitivity in response to NSC49L and TRAIL treatments. The results showed that indeed the overexpression of DR5 increased TRAIL-mediated sensitization of HT29-tet-DR5+dox cells than the HT29-tet-DR5 cells (Fig. 6B). The NSC49L treatment also sensitized the HT29-tet-DR5+dox cells, suggesting a link between NSC49L and TRAIL actions (Fig. 6C). However, the combination of TRAIL with NSC49L did not further affect the sensitization of these cells (Fig. 6C). This may be due to a saturating effect of NSC49L on the DR5 pathway that does not require TRAIL for further activation.

**Figure 6.**
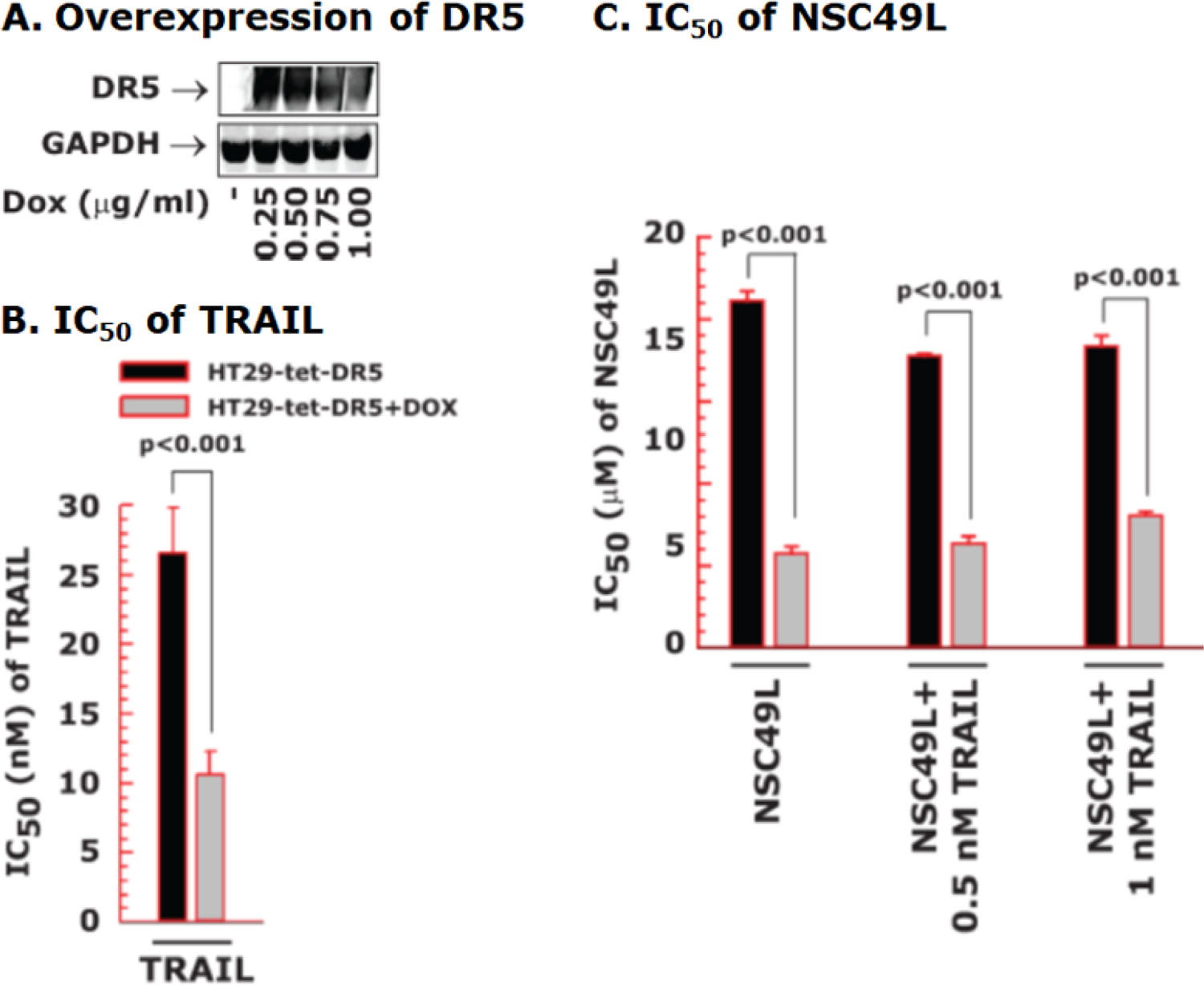
Overexpression of DR5 induces the sensitization of HT29 cells to NSC49L and TRAIL. One set of the HT29-tet-DR5 cells were treated with DOX (1 µg/ml) to induce the expression of DR5. Then, control and DR5-induced cells were treated with different concentrations of NSC49L and TRAIL either alone or in combination for 72 h. The cell survival was determined by the MTT-assay. **A,** Doxycycline-induced DR5 expression was verified by immunoblot. **B,** IC50 of TRAIL. **C,** IC50 of NSC49L in the presence of 0, 0.5 and 1 nM of TRAIL, respectively. Data are the mean ± SE of four different estimations.

### NSC49L and TRAIL treatments downregulate AKT1/mTOR/4EBP1 signaling in FOLFOX-HT29 cells

In these experiments, we determined a link between AKT1, mTOR and 4EBP1 in NSC49L- and TRAIL-treated FOLFOX-HT29 cells. The results showed that there was no effect of the NSC49L and TRAIL treatments on the total levels of AKT1, AKT2 and AKT3 in FOLFOX-HT29 cells; however, the AKT1(S473P) level was significantly reduced (Fig. 7A).

**Figure 7.**
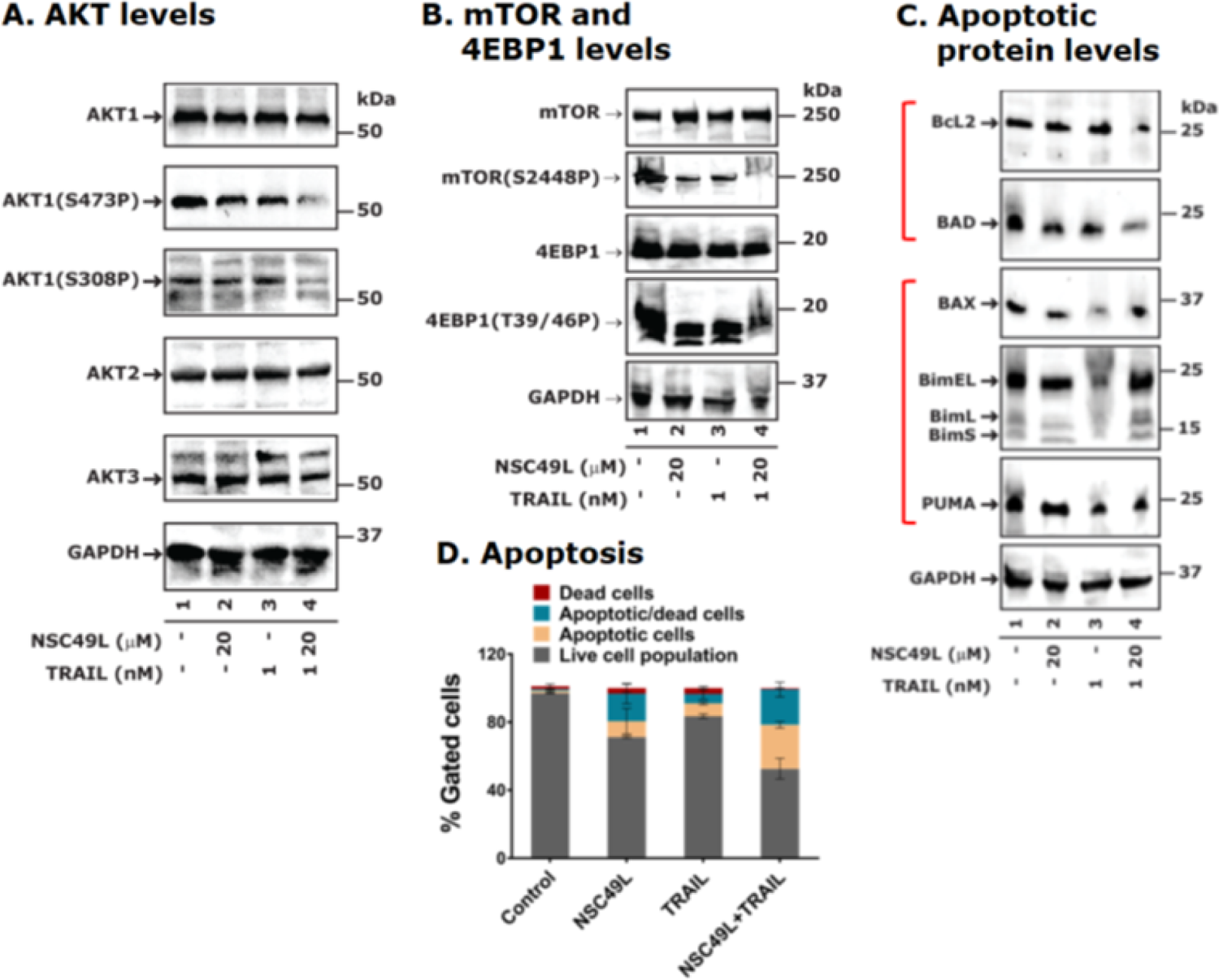
Western blot analysis of AKT/mTOR/4EBP1, apoptotic pathway proteins and apoptosis of FOLFOX-HT29 cells-treated with NSC49L and TRAIL either alone or in combination for 24 h. **A** and **B**, western blots demonstrating regulation of the AKT/mTOR/4EBP1 pathway by NSC49L and TRAIL in FOLFOX-HT29 cells. **C,** levels of apoptotic proteins. All of the blots are the representative of three experiments. **D,** caspase 3/7 activity. FOLFOX-HT29 cells were treated with NSC49L and TRAIL either alone or in combination for 24 h, and the extent of apoptosis was determined by measuring the activation of caspase 3/7 using a flow cytometer. The experiment was repeated twice. Data are the mean ± SE of triplicate results.

The AKT1(T308P) level was also reduced, although not as much (Fig. 7A). These results suggest the importance of AKT1(S473P) in the induction of FOLFOX-resistance in HT29 cells. Further, the phosphorylation of mTOR(S2448P) and its downstream target protein 4EBP1(T37/46P) was also decreased in NSC49L and TRAIL-treated FOLFOX-HT29 cells (Fig. 7B). From these results, we conclude that NSC49L and TRAIL treatment affect the AKT1/mTOR/4EBP1/p21-axis in FOLFOX-resistant CRC cells.

### NSC49L- and TRAIL treatment increased the apoptosis of FOLFOX-HT29 cells

We examined the expression levels of pro-apoptotic (Bcl2 and BAD) and anti-apoptotic (BAX, Bim and PUMA) proteins in FOLFOX-HT29 cells. The results showed a decrease in Bcl2 and BAD and an increase in Bax and Bim, but not PUMA, protein levels in FOLFOX-HT29 cells-treated with NSC49L and TRAIL (Fig. 7C), which may be sufficient to drive these cells to cell death. To further confirm that NSC49L and TRAIL treatment induces apoptosis in FOLFOX-HT29 cells, we measured caspase 3/7 activation, which are executioner caspases that mediate apoptotic cell death (McIlwain et al., 2013). The results showed increased apoptosis after NSC49L treatment as compared to control, which was further increased when it was combined with TRAIL (Fig. 7D).

TRAIL alone also caused apoptosis of FOLFOX-HT29 cells, but less than NSC49L alone (Fig. 7D). Similar results were also observed with FOLFOX-HCT116 cells (Fig. S4). These results suggest that combining NSC49L and TRAIL decreases the levels of the pro-apoptotic proteins, increases the levels of the anti-apoptotic proteins, and induces apoptosis in FOLFOX-resistant cells.

### NSC49L and TRAIL synergistically enhance the sensitization of p21-knockout CRC cells

Next, we determined the importance of p21 in the sensitization of CRC cells in response to NSC49L and TRAIL treatments. For these experiments, we used isogenic wild-type and p21-knockout HCT116 cell lines. We also overexpressed H6p21 (6 his-tagged p21) to reverse the NSC49L- and TRAIL-mediated sensitization effect observed with p21-knockout cells. The HCT116-p21^-/-^ cells were highly sensitive to TRAIL (IC50 of 0.54 ± 0.18 nM) (Fig. 8A), and the sensitivity was decreased after the overexpression of H6p21 (IC50 of 1.46 ± 0.34 nM) (Fig. 9A). Further, the sensitization of HCT116-p21^-/-^ cells drastically increased when the cells were treated together with NSC49L and TRAIL (Fig. 8B). These results suggested that the combination of NSC49L and TRAIL have a profound effect on the sensitization of HCT116-p21^-/-^ cells compared to HCT116-p21^+/+^ cells. To explore the possibility that the NSC49L and TRAIL treatments on HCT116-p21^-/-^ cells showed a synergy, we determined the combination index (CI) by applying the Chou-Talalay method (Chou, 2010). The CI calculated by the Chou-Talalay approach provides a quantitative definition for synergism (CI < 1), additive effects (CI= 1), or antagonism (CI > 1). Based on this method, we performed MTT-cell viability assays with HCT116-p21^-/-^ cells using increasing concentrations of NSC49L and TRAIL, where the increment of both concentrations was kept at a 1:1 ratio. The data analysis confirmed that the combined treatment with NSC49L and TRAIL has a synergistic effect in the HCT116-p21^-/-^ cells (Fig. 8C). This effect was reversed in the HCT116-p21^-/-^/H6p21 cells that overexpress p21. The HCT116-p21^-/-^/H6p21 cells showed more resistance to NSC49L and TRAIL than HCT116-p21^-/-^ cells (Fig. 9A). The IC50 of NSC49L in the presence of TRAIL became 5.5-fold higher in HCT116-p21^-/-^/H6p21 cells than the IC50 of HCT116-p21^-/-^ cells (Fig. 9B). Similarly, the IC50 of TRAIL in the presence of NSC49L became 10.67-fold higher in HCT116-p21^-/-^/H6p21 cells than the IC50 of HCT116-p21^-/-^ cells (Fig. 9C). These results support our hypothesis that the downregulation of p21 is critical for NSC49L- and TRAIL-mediated sensitization of CRC cells.

**Figure 8.**
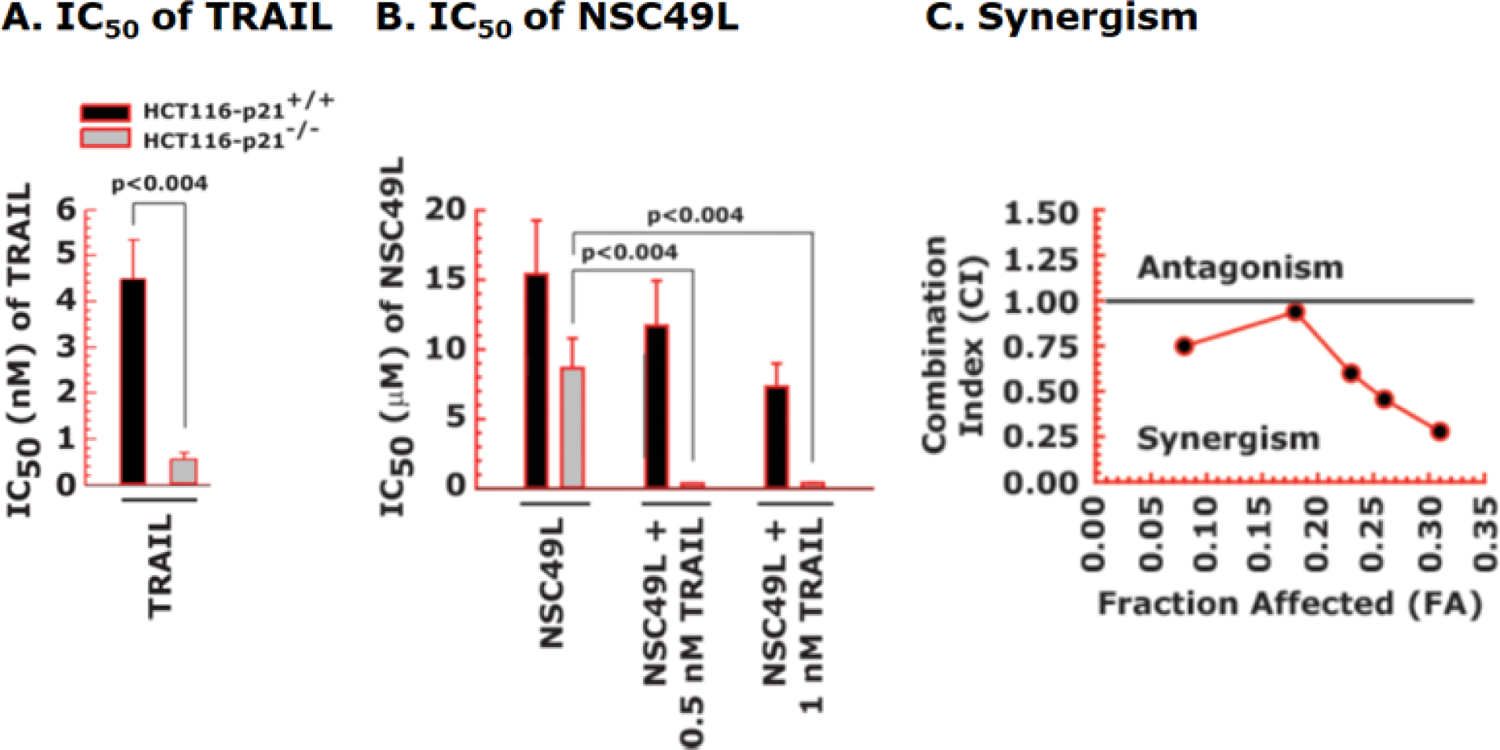
Synergistic sensitization of p21-knockout cells in response to NSC49L and TRAIL treatment. HCT116-p21^+/+^ and HCT116-p21^-/-^ cells were treated with different concentrations of NSC49L and TRAIL either alone or in combination for 72 h and the cell survival was determined by the MTT-assay. **A** and **B,** the IC50 of TRAIL and NSC49L cells, respectively. The IC50 is derived from the survival curves. **C,** The synergistic effect of NSC49L and TRAIL with HCT116-p21^-/-^ cells. Data are the mean ± SE of four estimations.

**Figure 9.**
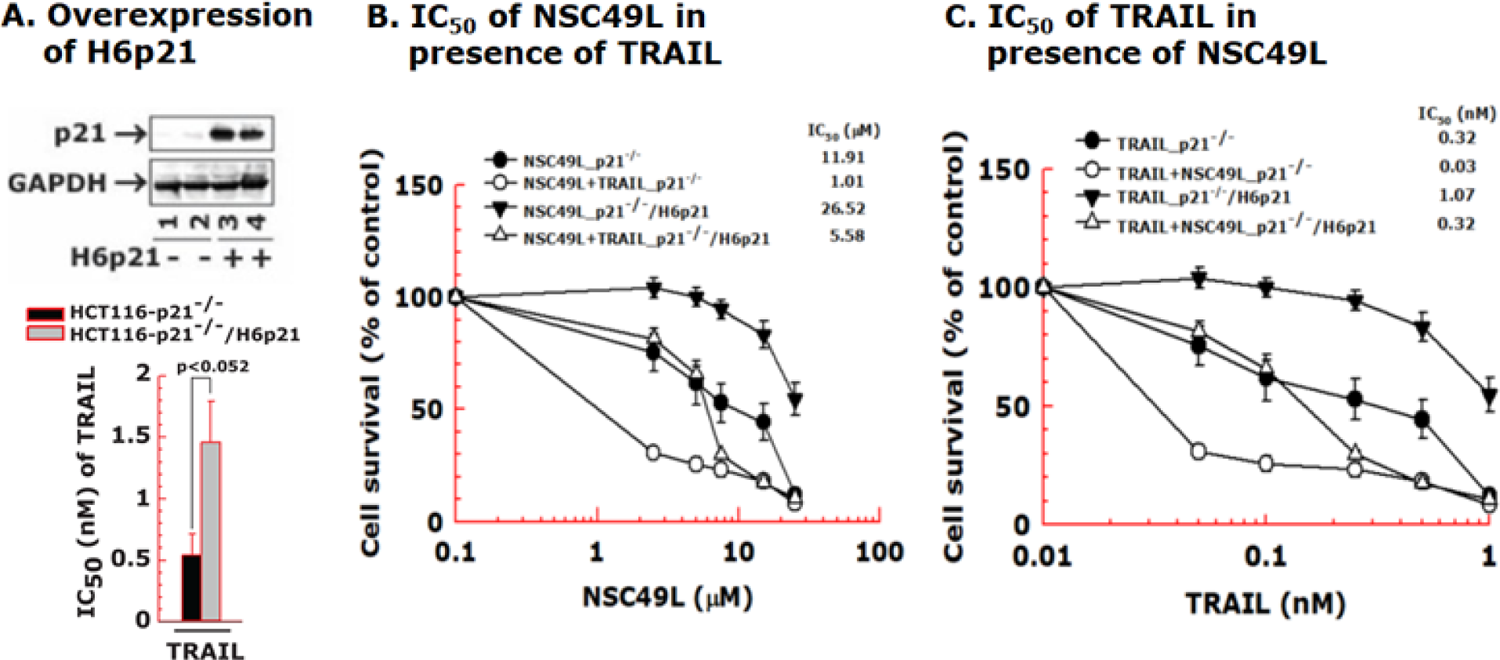
Overexpression of H6p21 decreases the sensitization of p21-knockout cells in response to NSC49L and TRAIL treatment. The HCT116-p21^-/-^ and HCT116-p21^-/-^/H6p21 cells were treated with different concentrations of NSC49L and TRAIL either alone or in combination for 72 h and the cell viability was determined by the MTT-assay. **A,** western blot showing the overexpression of H6p21. **B,** the IC50 of NSC49L in combination with TRAIL. **C,** IC50 of TRAIL in combination with NSC49L. Different concentrations of NSC49L (0, 2.5, 5, 7.5, 15 and 25 µM) and TRAIL (0, 0.05, 0.1, 0.25, 0.5 and 1.0 nM) were used in the synergy experiments. Data are the mean ± SE of four estimations.

### NSC49L treatment increases *CDKN1A* mRNA levels and does not affect p21 protein degradation

The increased level of p21 in FOLFOX-HT29 cells is independent of p53 because the *TP53* gene is mutated in these cells (Rodrigues et al., 1990). While the mutant *TP53* expression is higher in HT29 cells, it is drastically reduced in FOLFOX-HT29 cells, possibly due to transcriptional silencing of *TP53* (Fig. 10A, left). This supports the loss of the p53 protein level shown in Figure 5E. Now, the question arises whether the decreased level of p21 in FOLFOX-HT29 cells after NSC49L treatment is secondary to decreased transcription or increased protein degradation. First, we performed qRT-PCR to examine whether NSC49L-induced low levels of p21 protein is due to decreased *CDKN1A* gene transcription. We treated HT29 and FOLFOX-HT29 cells with 20 µM of NSC49L for 16 h, then, determined *CDKN1A* mRNA levels by qRT-PCR. The results showed an increased level of *CDKN1A* mRNA in FOLFOX-HT29 cells that was not further altered after NSC49L treatment (Fig. 10A, right).

**Figure 10.**
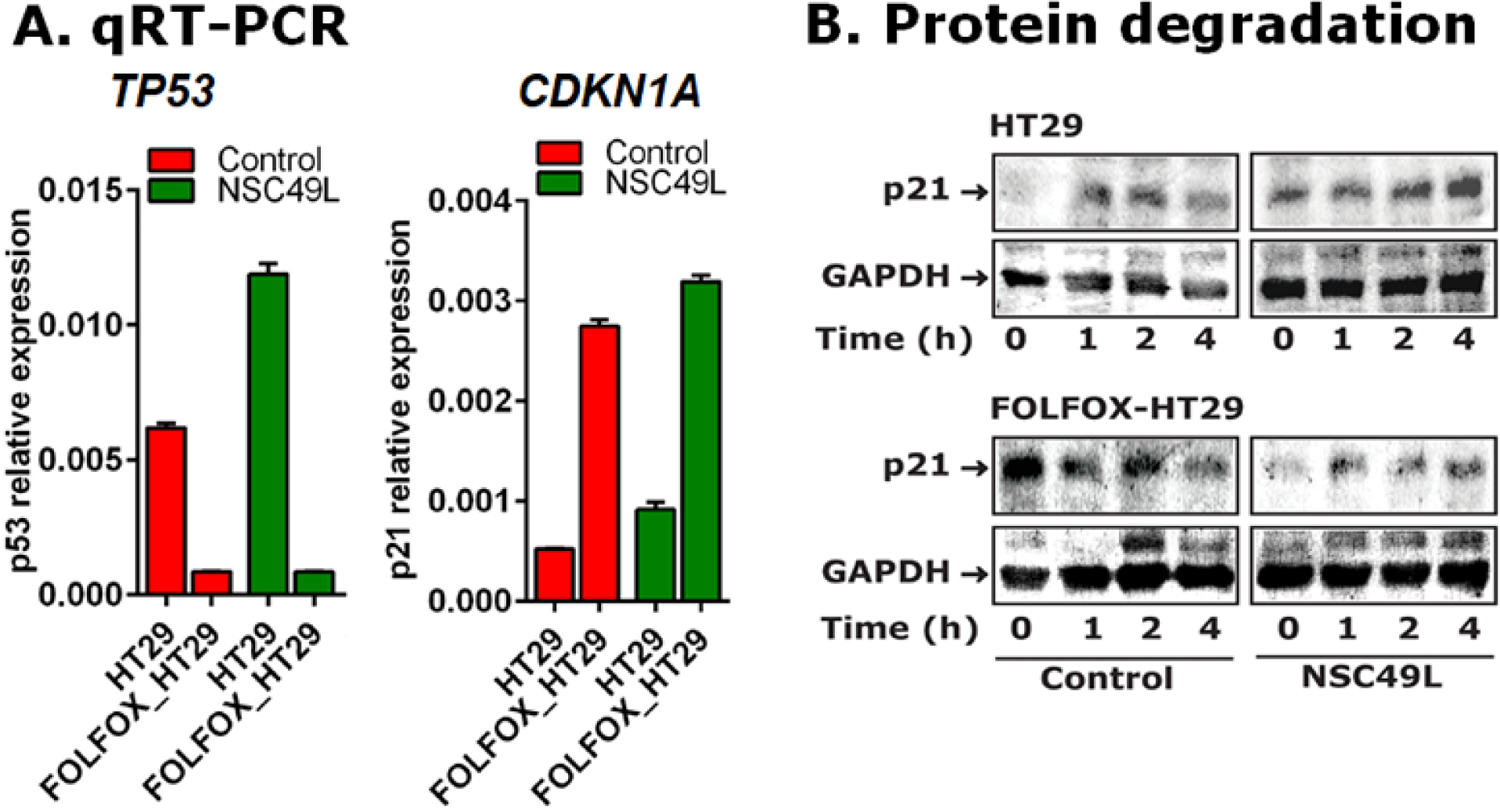
Effect of NSC49L and TRAIL treatments on the *TP53* and *CDKN1A* mRNA levels and protein kinetics in HT29 and FOLFOX-HT29 cells. A, cells were treated with NSC49L (20 µM) and TRAIL (1 nM) either alone or in combination for 16 h. Total RNA was isolated and used for qRT-PCR. Data are the mean ± SE of four estimations. **B,** kinetics of p21 protein stability after CHX treatment. Cells were treated with NSC49L (20 µM) and TRAIL (1 nM) either alone or in combination for 24 h followed by the treatment with CHX (10 µg/ml) for different periods. The p21 protein level was determined by western blot analysis. Blots shown are the representative of two experiments.

Similarly, we observed no effect of NSC49L treatment on *CDKN1A* levels in FOLFOX-HCT116 cells (Fig. S5). These results indicate that NSC49L does not change *CDKN1A* gene expression in FOLFOX-HT29 and FOLFOX-HCT116 cells; however, the increased *CDKN1A* mRNA levels could be due to acquired *CDKN1A* gene expression during the development of FOLFOX-induced resistance.

Next, we examined whether the decreased p21 protein level in FOLFOX-HT29 cells after NSC49L treatment was secondary to protein degradation. We treated HT29 and FOLFOX-HT29 cell lines with 20 µM NSC49L for 24 h. Then, we blocked nascent protein synthesis by treating the cells with 10 µg/ml of cycloheximide (CHX), which interferes with the translocation step in protein synthesis. Since p21 protein has a short half-life (Beuvink et al., 2005), we followed its levels up to 4 h. The results showed no apparent difference in the kinetics of p21 protein degradation in FOLFOX-HT29 cells. Similar results were observed with HT29 cells as well (Fig. 10B). These results suggest that NSC49L treatment-mediated decrease of p21 protein levels in FOLFOX-HT29 cells is not due to decreased mRNA levels or increased protein degradation, but it could be due to a decrease in protein translation.

### The rate of p21 protein synthesis is reduced by NSC49L treatment in FOLFOX-HT29 cells

It is known that hypophosphorylated 4EBP1 binds to eIF4E and prevents it from interacting with eIF4G to promote ribosome recruitment to mRNAs. Hence, this can repress the initiation of mRNA translation. Once the phosphorylated 4EBP1(T37/46P) dissociates from eIF4E, it can then allow for the formation of the eIF4F complex and the initiation of cap-dependent translation (Lin et al., 1994). As discussed above, we observed that the increased level of 4EBP1(T37/46P) in FOLFOX-HT29 cells decreased after the treatment with NSC49L (Fig. 7B). These results suggested that the decreased level of 4EBP1(T37/46P) may provide an opportunity to increase the interaction of hypophosphorylated 4EBP1 with eIF4E in these cells that may occur after treatment with NSC49L. To test this possibility, we performed immunohistochemistry (IHC) experiments with FOLFOX-HT29 cells. Since untreated cells have higher levels of 4EBP1(T37/46P), we did not observe colocalization of 4EBP1(T37/46P) and eIF4E. However, as the level of 4EBP1(T37/46P) decreased after NSC49L treatment (Fig. 7B), we observed an increased colocalization of hypophosphorylated 4EBP1 with eIF4E (appearance of yellow color) (Fig. 11A). In HT29 cells, we observed very poor colocalization of 4EBP1 with eIF4E after NSC49L treatment (Fig. S6). This correlates with the low level of p21 in these cells due to the presence of mutant p53, which could not regulate *CDKN1A* gene expression (Fig. 5E). These results suggest that the NSC49L treatment decreases the levels of 4EBP1(T37/46P) and increases the colocalization of 4EBP1 with eIF4E, which may block eIF4F complex assembly, and hence decrease the translation of target protein(s), such as p21.

**Figure 11.**
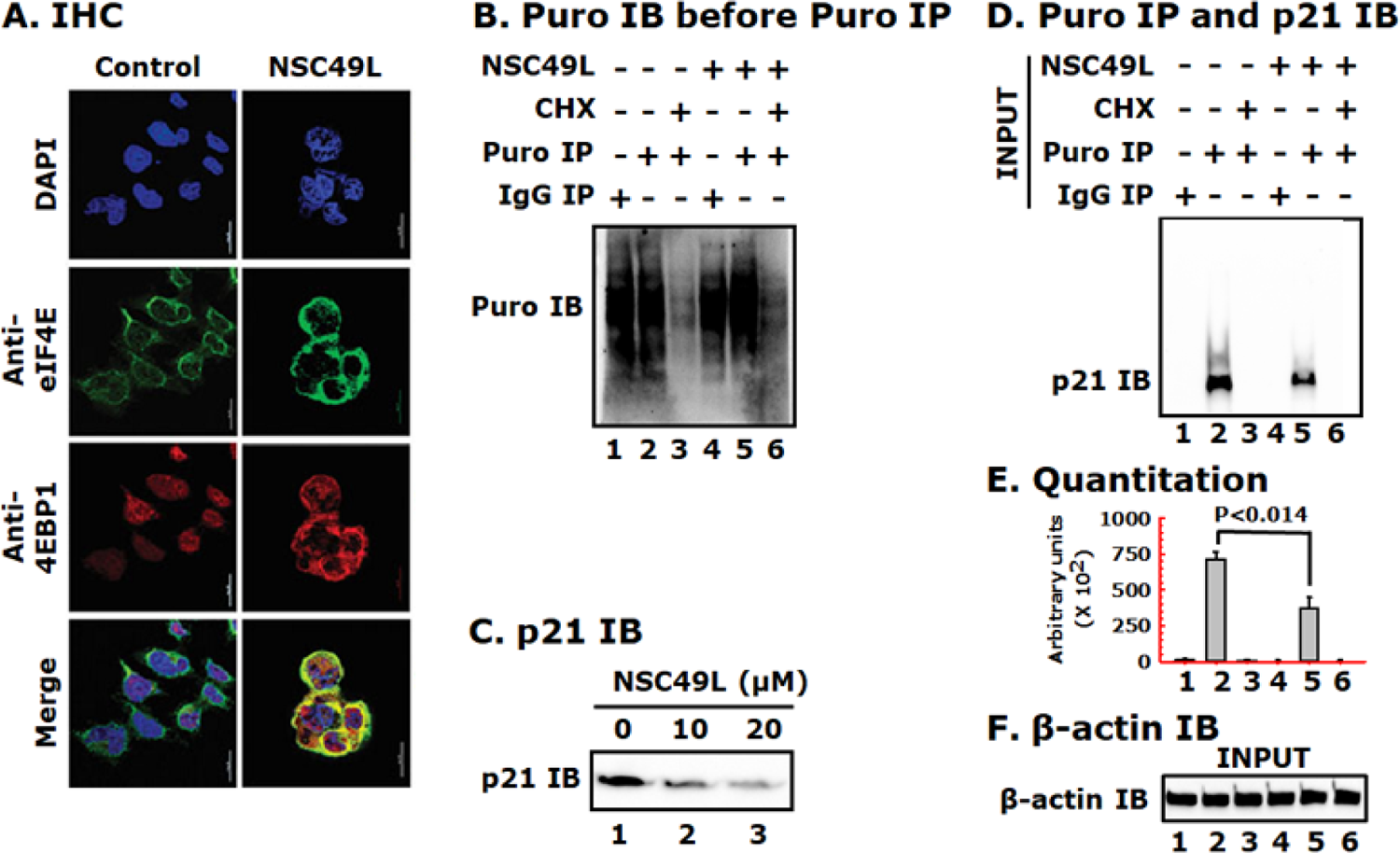
NSC49L treatment increase the colocalization of eIF4E with unphosphorylated 4EBP1 and inhibits p21 protein synthesis in FOLFOX-HT29 cells. A, **IHC analysis.** FOLFOX-HT29 cells were treated with 20 µM of NSC49L for 24 h. After treatment, the colocalization of eIF4E and 4EBP1 was determined by IHC. Yellow color in merge column shows the colocalization of these two proteins. Images were captured with a Nikon A1RMPsi-STORM4.0 confocal microscope (Melville, NY) at 60X magnification. **B,** global protein synthesis. FOLFOX HT29 cells were treated as above with 20 µM of NSC49L for 24 h. Cells were treated with 10 µg/ml of puromycin or puromycin plus 10 µg/ml of CHX for 15 min. Cells were washed and incubated in fresh medium without drugs for additional 2 h. Samples were blotted with anti-puromycin antibody. **C,** validate the experimental conditions, a concentration dependent effect on p21 expression was performed. Cells were mock treated (control) or treated with 10 µM and 20 µM of NSC49L for 24 h and p21 level was determined. **D,** evaluate the effect of NSC49L on the regulation of p21 biosynthesis, we set up the experiment as in B. To determine the incorporation of puromycin into nascent p21, the pre-immunoprecipitated lysate was subjected to anti-puromycin immunoblot. **E,** quantitative analysis of the blot shown in panel D, which is presented as the ImageJ arbitrary units. Data are Mean ± SE of three experiments. **F,** β-actin was used as an internal control for equal loading.

Then, to examine the effect of NSC49L on p21 protein synthesis, we used the puromycin-based surface sensing of translation (SUnSET) assay protocol for labeling of nascent proteins (Schmidt et al., 2009). Puromycin functions as a structural analog of aminoacyl tRNAs and leads to the release of unfinished nascent polypeptide chains during the ribosomal elongation cycle and induces premature termination and subsequent drop-off of the ribosome from the mRNA. The relative rate of puromycylated nascent proteins can be detected by anti-puromycin antibody in control and drug-treated cells (Iwasaki and Ingolia, 2017). Here, we treated FOLFOX-HT29 cells with 20 µM of NSC49L for 24 h. Then, changed the medium and followed a 2 h chase with 10 µg/ml of puromycin, prepared whole cell lysates and performed immunoprecipitation (IP) and immunoblotting (IB). In the case of cycloheximide (CHX), after 24 h of NSC49L treatment, we treated the cells with 10 µg/ml of CHX for 1 h, followed by 2 h of chase with puromycin. In contrast to puromycin, CHX inhibits protein synthesis by binding exclusively of cytoplasmic (80S) ribosomes of eukaryotes (Stocklein and Piepersberg, 1980). Pre-treatment with cycloheximide can limit puromycin incorporation by blocking peptidyl-tRNA transition from the A-site (Hobden and Cundliffe, 1978). First, we determined the effect of NSC49L on global protein synthesis in FOLFOX-HT29 cells. The results showed that global protein synthesis was unaffected by NSC49L treatment, while it was blocked by the CHX treatment (Fig. 11B, compare lane 2 with 5 and 3 with 6, respectively). Treatment with NSC49L caused a dose-dependent decrease in p21 protein levels (Fig. 11C), confirming the validity of the experimental conditions. Next, we determined the effect of NSC49L treatment on the rate of nascent p21 protein synthesis. We found a significantly decreased rate of nascent p21 synthesis in FOLFOX-HT29 cells after NSC49L treatment as compared to control (Fig. 11D and E, compare lane 2 with 5). CHX treatment completely blocked p21 synthesis (Fig. 11D and E, compare lane 3 with 6). Data in lanes 1 and 4 show the IgG control. The β-actin protein level indicates that the same amounts of whole cell lysate were used in IP experiments (Fig. 11F). These results thus suggest that NSC49L reduces the rate of p21 protein translation in FOLFOX-HT29 cells linked with the PP2A/AKT1/mTOR/4EBP1-axis.

### The TRAIL pathway interacts with the NSC49L pathway at the level of procaspase 3 and p21

As discussed above, we consistently observed a profound effect of NSC49L when applied in combination with TRAIL on increased cytotoxicity, increased c-Caspase 8, c-Caspase 3, c-PARP1, and decreased p21 protein levels in FOLFOX-resistant CRCs. Since TRAIL is not expected to affect PP2A activity, its coordination with NSC49L may occur at the p21 level. In previous studies, the interaction of p21 with procaspase 3, and thus the blockade of Fas-induced apoptosis has been shown (Suzuki et al., 1998; Suzuki et al., 1999). There is also evidence that caspase 3 cleaves p21, thereby, inducing apoptosis in various cell types (Jin et al., 2000). Based on these findings, we predicted that TRAIL might induce procaspase 3 cleavage into caspase 3 once free from p21 after downregulation through the NSC49L pathway. Then, the active caspase 3, through a feedback mechanism, may further cleave p21 to enhance the rate of apoptosis. If this is true, then inhibition of caspase 3 activity may block NSC49L and TRAIL-induced sensitization of FOLFOX-HT29 cells and the cleavage of procaspase-3 and PARP1.

To test this possibility, we first examined the localization of p21 and procaspase-3 in response to NSC49L and TRAIL using immunofluorescence microscopy. We treated HT29 and FOLFOX-HT29 cells grown on cover slips with 20 µM of NSC49L and 1 nM of TRAIL either alone or in combination for 24 h. Cells were processed for IHC using anti-p21 and anti-procaspase antibodies with corresponding AlexaFluor-488 (green) and AlexaFluor-568 (red) secondary antibodies. Images were captured with a Leica DM IRBE microscope (Hawthorn, NY) at 20X magnification. Since p21 levels are very low in HT29 cells, the colocalization of p21 and procaspase 3 was also poor; however, after NSC49L and TRAIL treatment it showed reduced levels and colocalization (Fig. S7A). On the other hand, we observed a higher level of p21 and its colocalization with procaspase in FOLFOX-HT29 cells, which was drastically reduced after the NSC49L but not the TRAIL treatment (Fig. S7B). The combination treatment of NSC49L and TRAIL even further reduced the levels and the colocalization of p21 and procaspase 3 (Fig. S7B). These results suggest that p21 may interact with procaspase 3 and keep it in its inactive state in FOLFOX-HT29 cells. Procaspase 3 may become free and get cleaved to active caspase 3 after NSC49L-mediated p21 reduction in these cells.

Then, we examined whether the caspase 3 specific inhibitor, ZDEVD, will block NSC49- and TRAIL-induced sensitization of FOLFOX-HT29 cells. For these experiments, we pretreated FOLFOX-HT29 cells with 20 µM of ZDEVD for 3 h, followed by the treatment with NSC49L and TRAIL either alone or in combination for an additional 72 h. We performed MTT-cell viability assays and determined the IC50 of the drugs. Here, we have shown the sensitization effect of the drugs as IC50. Pretreatment with ZDEVD significantly increased the IC50 of TRAIL as compared to control, while it did not affect the IC50 of NSC49L (Fig.12A and B, respectively). However, pretreatment with ZDEVD significantly increased the IC50 of NSC49L when used in combination with TRAIL (Fig. 12B). These results suggest that ZDEVD is acting through the TRAIL pathway, and the NSC49L pathway is coordinating with the TRAIL pathway. Next, we examined whether the coordination between the TRAIL and NSC49L pathways are at the procaspase 3 level. We used the same protocol as above, except that the experiments were terminated after 24 h. Since we wanted to examine the effect of ZDEVD on TRAIL-induced effects, we treated FOLFOX-HT29 cells with 1 nM of TRAIL with or without 10 µM of ZDEVD and 20 µM of NSC49L. Western blot results showed no appreciable change in the level of DR5 after combination treatment of TRAIL and NSC49L as compared to TRAIL alone treated cells (Fig. 12C, compare lane 1 with 2). DR5 levels also remained unchanged after TRAIL and NSC49L treatment in the ZDEVD-pretreated cells (Fig. 12C, compare lane 3 with 4). On the other hand, p21 levels were decreased in the TRAIL and NSC49L combination treatments as compared to TRAIL alone treated cells (Fig. 12C, compare lane 1 with 2), which was restored in the ZDEVD-pretreated cells (Fig. 12C, compare lane 3 with 4). Further, the c-Caspase 8, c-Caspase 3 and c-PARP1 levels were increased, and the procaspase 3 level was decreased in (Fig. 12C, compare lane 1 with 2). Interestingly, in the ZDEVD-pretreated cells, the combined effects of TRAIL and NSC49L on c-Caspase 8, c-Caspase 3 and c-PARP1, and procaspase 3 levels were reversed (Fig. 12C, compare lane 3 with 4). Taken together, these results establish a relationship between p21 and caspase 3 in the regulation of NSC49L- and TRAIL-induced cytotoxic activities in FOLFOX-HT29 cells.

**Figure 12.**
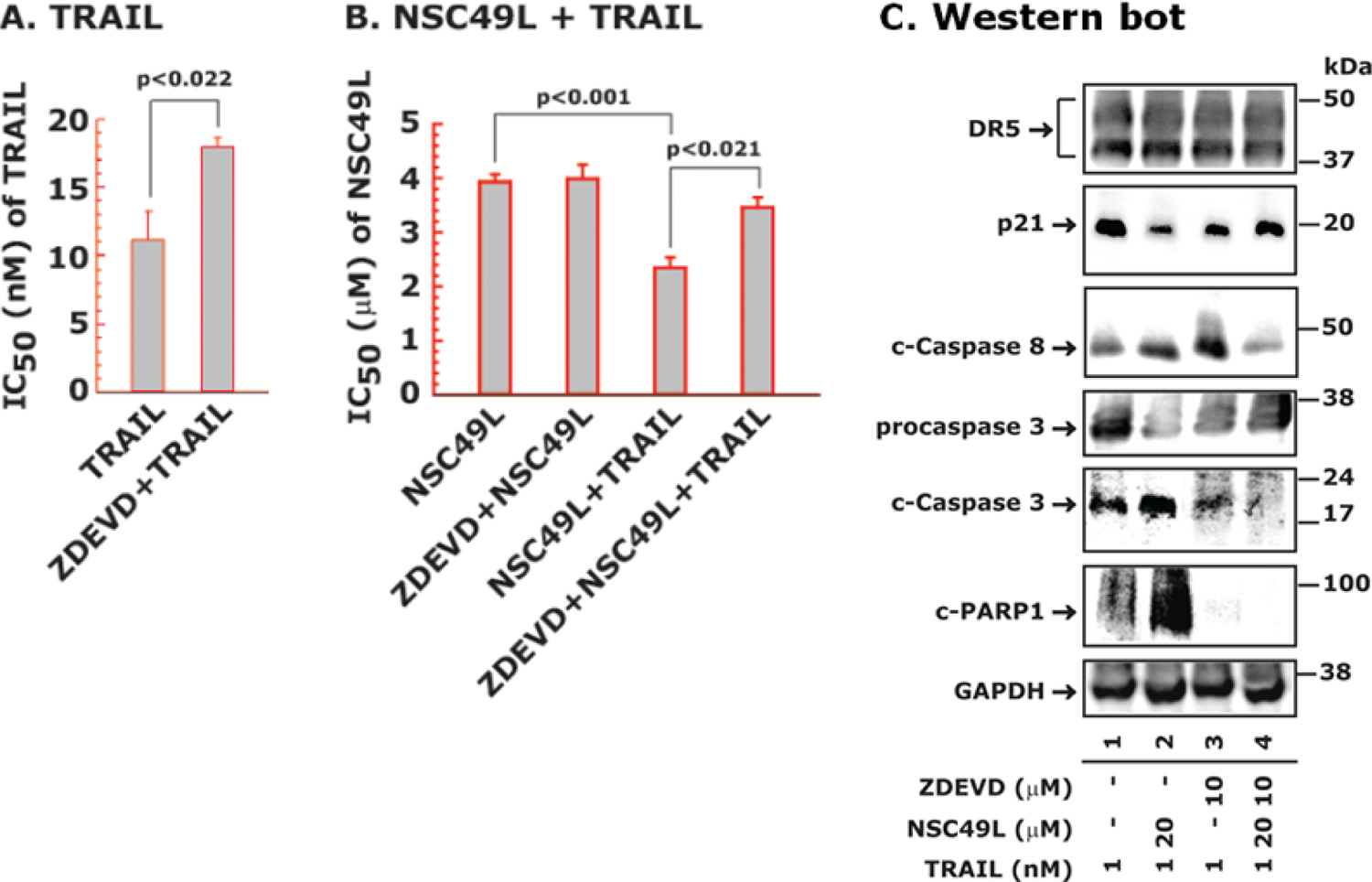
Caspase 3 inhibitor ZDEVD reverses the cytotoxic effect of NSC49L and TRAIL in FOLFOX-HT29 cells. Cells were pretreated with 20 µM ZDEVD for 3 h followed by the treatment with different concentrations of NSC49L and TRAIL either alone or in combination for an additional 72 h. Cell viability was determined using the MTT-assay. **A,** the IC50 of TRAIL. **B,** the IC50 of NSC49L in combination of TRAIL. Data are the mean ± SE of four estimations. **C,** western bots. For these experiments, after pretreatment with ZDEVD, the treatment with NSC49L and TRAIL either alone or in combination was carried out for 24 h. Blots are the representative of two different experiments.

## DISCUSSION

Access to TCGA data through the Genomic Data Commons Data Portal has provided an enormous opportunity to analyze important genes and pathways by which they can be implicated in developing precision medicine approaches for the diagnosis and treatment of cancer (Jensen et al., 2017). First, we found a decreased level of *PPP2C* in CRC tumors compared to normal tissues, and this pattern was linked with the progression of the tumor through different stages.

Then, we analyzed CPTAC data and found a similar trend at the protein level. The low expression of *PPP2CB* was linked with poor prognosis of CRC patients. Second, we used a meta-analysis method to analyze three microarray datasets (accession number (GSE19860, GSE28702 and GSE72970)) of FOLFOX-responsive and non-responsive CRC tumors (Lin et al., 2018), and established a link between *PPP2C* and *AKT1*, *mTOR*, *4EBP1* and *CDKN1A*. We found a decreased level of *PPP2CA/B* in FOLFOX-non-responsive compared with responsive tumors. Further, the expression levels of *AKT1*, *mTOR*, *4EBP1* and *CDKN1A* were increased in FOLFOX-non-responsive compared with responsive tumors. These results provided clear evidence that upregulation of PP2A and downregulation of the AKT1, mTOR, 4EBP1 and p21 levels can be beneficial for the management of CRC patients.

In the present study, we pursued the above concept and identified a novel PP2A agonist, NSC49L, and described the molecular mechanisms by which it downregulates AKT/mTOR/4EBP1 pathway-dependent p21 translation and induces TRAIL/caspase-3-dependent apoptosis of FOLFOX-resistant CRC cells. Decreased levels of PP2AC (Yong et al., 2018) and increased levels of phospho-AKT (Li, 2018), phospho-mTOR (Melling et al., 2015), phospho-4EBP1 (Malinowsky et al., 2014) and TRAIL (Zhang and Fang, 2005) have been implicated as drug-resistance factors for metastatic CRC patients. We have identified p21 as an acquired resistance factor that is observed in HCT116 and HT29 cells during FOLFOX treatment. P21 levels were increased in both FOLFOX-HCT116 and FOLFOX-HT29 cell lines in a p53-independent manner. In fact, we observed loss of p53 expression in both FOLFOX-HCT116 and FOLFOX-HT29 cell lines. This is the first report showing the upregulation of p21 and loss of p53 in FOLFOX-resistant CRC cells. The role of p21 is well-established in protecting CRC cells against a variety of stress stimuli, including exposure to radiation and chemotherapy, such as 5-FU (Bene and Chambers, 2009; Gorospe et al., 1996; Mahyar-Roemer and Roemer, 2001; Maiuthed et al., 2018; Sharma et al., 2005; Tian et al., 2000). Its upregulation correlates positively with tumor grade, invasiveness and aggressiveness, and is considered as a poor prognostic indicator (Abbas and Dutta, 2009). An anti-apoptotic and enhanced cell survival role of p21 is implicated by inhibiting initiator caspases directed by the TRAIL/DR4 pathway (Xu and El-Deiry, 2000). Considering the anti-apoptotic role of p21, it is therefore, emerging as a therapeutic target for certain cancers (Weiss, 2003). While in the current study we focused on p21 as a target of PP2A agonist-induced sensitization of FOLFOX-resistant CRC cells, the mechanism of the loss of p53 expression is still unclear.

By using inhibitors, a link between AKT1, mTOR and 4EBP1 has been observed (Wang et al., 2020; Wang and Zhang, 2014); however, their efficacy in sensitizing FOLFOX-resistant CRC cells is not clear. Further, the link between PP2A and the AKT1/mTOR/4EBP1-axis and with TRAIL is also obscure. Since PP2A dephosphorylates AKT1 and induces sensitization of cancer cells, it is desirable to develop a PP2A activator. In the past, several PP2A agonists have been developed that showed a significant therapeutic efficacy when used in combination with oncogenic kinase inhibitors, DNA damaging agents and radiation (Mazhar et al., 2019). These agonists modulate the activity of PP2A by interacting with its scaffolding (*PPP2R1A*, PR65α) and regulatory (*PPP2R2A*, B55α) subunits, or by removing the antagonistic effects of the PP2A regulatory subunits (Mazhar et al., 2019; McClinch et al., 2018; Sangodkar et al., 2017).

Recently, phenothiazine derivatives have been identified as PP2A agonists that regulate enzymes containing the *PPP2R5E* (B56ε) regulatory subunit (Morita et al., 2020). However, the targeting by agonists to PP2A’s catalytic (C) subunit is largely unknown. Further, these agonists have not been tested against FOLFOX-resistant CRC cells, and show a poor safety profile as well. On the other hand, our PP2A agonist, NSC49L, is specific to its catalytic subunit, has a safer profile, and sensitizes FOLFOX-resistant CRC cells *in vitro* (present data) and *in vivo* patient-derived xenograft models of CRC stem-like cell tumors (Narayan et al., 2017). NSC49L inhibits the AKT1/mTOR/4EBP-axis and interrupts p21 translation. The reduced level of p21 then links with the TRAIL pathway, thereby, inducing caspase 3 activation and sensitization of FOLFOX-resistant CRC cells.

In previous studies, cap-dependent translational control has been implicated as a therapeutic target and biomarker for cancer (Vaklavas et al., 2017). Translational control of p21 has been reported for mTOR inhibitors (Beuvink et al., 2005), nutrient stress (Lehman et al., 2015) and UVB-irradiation (Collier et al., 2018). The first step in the cap-dependent translation initiation is the assembly of the trimolecular cap-binding complex, eIF4F, at the 5’UTR of the mRNA bound to the ribosome (Bhat et al., 2015). EIF4F consists of three subunits: eIF4E, the cap binding protein; eIF4A, a bidirectional ATP-dependent RNA helicase; and eIF4G, a modular scaffolding protein that binds eIF4E, eIF4A, eIF3 and poly(A)-binding protein (PABP). A coordinated interaction between eIF4G, eIF4E and PABP brings about mRNA circularization and promotes translation initiation (Kahvejian et al., 2005). The colocalization of eIF4E and eIF4G at the 5′-m^7^GpppN mRNA-cap is a critical step in translational control. The binding of 4EBP1 to the same region of eIF4E where eIF4G binds blocks translation (Bhat et al., 2015). Phosphorylated 4EBP1 dissociates from the complex and allows eIF4E to form an active translation initiation complex (Lin et al., 1994; Pause et al., 1994). The results of our studies supported this hypothesis. The data showed an increased colocalization of 4EBP1 and eIF4E with a decreased p21 translation in FOLFOX-resistant cells. Since *CDKN1A* mRNA levels were increased in the FOLFOX-resistant cells, the possibility of *CDKN1A* sequestration into stress granules (SGs) is likely. SGs are cytoplasmic multimeric RNA bodies formed in response to anticancer drugs. The sequestered *CDKN1A* may become unavailable for protein translation in the FOLFOX-resistant cells after the NSC49L treatment. In recent years, the involvement of AKT/mTOR/4EBP1 has been described in the formation of SGs in response to anticancer drugs (Fournier et al., 2013).

The formation of SGs is known to inhibit cap-dependent translation of p21 by the stabilization of *CDKN1A* mRNA in SGs (Gareau et al., 2011). The proteasome inhibitor, MG132, causes trapping of *CDKN1A* mRNA in SGs resulting in the inhibition of p21 translation (Lian and Gallouzi, 2009). We have not examined the involvement of SGs in NSC49L-mediated regulation of p21 translation. Nonetheless, our results clearly suggest that NSC49L affects selective inhibition of p21 translation, instead the global inhibition of protein translation, and thus can be a more therapeutically desirable agent than the ones affecting global protein synthesis. In other studies, rapamycin-dependent selective inhibition of mitogen-induced p21 translation (Gaben et al., 2004), as well as a global inhibition by mTOR inhibitor RAD001 (Beuvink et al., 2005) has been reported. The global inhibition of translation may preferentially affect cancer cells because the cancer cells have higher rates of translation than normal cells (Heys et al., 1991; Mills et al., 2008).

There is huge clinical potential for TRAIL in cancer therapy due to its lack of toxicity to normal cells, but high toxicity to cancer cells (Deng and Shah, 2020). However, about 50% of cancer cells develop resistance to TRAIL (Walczak et al., 1999), thus limiting its therapeutic efficacy. Currently, multiple means, including the use of genotoxic agents, epigenetic modulators, synthetic small molecules, or immunotherapy either alone or in combination have been implicated to circumvent TRAIL resistance and to induce apoptosis (Thorburn et al., 2008).

TRAIL-mediated resistance can occur through two general mechanisms. First, through anti-apoptotic mechanisms, such as the reduced expression or epigenetic silencing of caspase 8 and increased expression of caspase inhibitors XIAP, cIAP2 and Mcl-1, or overexpression of anti-apoptotic protein Bcl2. Second, TRAIL resistance can result from decreased expression or cell surface localization of the TRAIL receptors DR4/DR5 or increased expression of DR4/DR5 inhibitors Fas-associated death domain-like interleukin-1β-converting enzyme-inhibitory protein (FLIP) and decoy receptor 1 and 2 (DcR1/DcR2) (Thorburn et al., 2008). In the present study, we have also observed that decreased expression of DR5 in HT29 caused TRAIL resistance, which was reversed after DR5 overexpression.

In distinct cancer types, different mechanisms of TRAIL resistance may be operative. However, in FOLFOX-resistant CRC cells we show p21 as a major TRAIL resistance factor. This is consistent with our results showing a synergistic effect of NSC49L and TRAIL on the sensitization of *CDKN1A* (*p21*)-knockdown CRC cells that is reversed after p21-overexpression. TRAIL-induced DR5/caspase 8 signaling is active in FOLFOX-resistant cells but becomes trapped at the procaspase 3 step, since p21 bound procaspase 3 is not a suitable substrate for caspase 8. Then, the trapped procaspase 3 is not processed into caspase 3, resulting in the onset of apoptosis and induction of TRAIL resistance (Fig. 12). After NSC49L treatment, PP2A activity was induced, which downregulated AKT1/mTOR/4EBP1 signaling and p21 translation in FOLFOX-resistant CRC cells. The reduced p21 level then relieved procaspase 3, which was then cleaved into caspase 3 through the TRAIL-induced DR5/caspase 8 pathway. Active caspase 3 then induced sensitization of FOLFOX-resistant CRC cells. Furthermore, active caspase 3 may through a feedback mechanism increase the cleavage of p21 to further potentiate the effect of NSC49L and TRAIL on the sensitization of these cells (Fig. 12). A feedback mechanism of caspase 3-mediated cleavage of p21 is consistent with previous findings (Jin et al., 2000; Zhang et al., 1999). Thus, these preclinical studies provide a mechanistic explanation for how the novel PP2A agonist NSC49L can modulate the PP2A/AKT1/mTOR/4EBP1-axis and enhance the therapeutic efficacy of TRAIL by targeting the downregulation of p21 translation and caspase 3 activation in FOLFOX-resistant CRC cells.

## SIGNIFICANCE

Despite the improved response rates of CRC to the combination of FOLFOX, these tumors quickly acquire resistance to FOLFOX. CRCs also become resistant to TRAIL which is a physiological means of killing CRC cells and sparing normal cells. PP2A is linked with cancer development and drug resistance by modulating the phosphorylation status of AKT1, mTOR and 4EBP1. We have shown that the AKT1/mTOR/4EBP1 axis in FOLFOX-resistant CRC cells selectively activates the translation of p21, which is linked with drug-resistance. Thus, activation of PP2A and downregulation of the AKT1/mTOR/4EBP1 axis and p21 translation may be a viable strategy for inducing sensitization of FOLFOX-resistant CRC cells. We have identified a novel PP2A activator, NSC49L, that decreases AKT1/mTOR/4EBP1 activity and the level of p21 in FOLFOX-resistant cells. We have described a link between p21 and procaspase 3 and TRAIL-resistance, and how a PP2A activator decreases p21 translation and activation of caspase 3-mediated sensitization of FOLFOX-resistant CRC cells. These studies will facilitate the development of a novel and clinically relevant PP2A activator and the identification of p21 as a novel downstream target involved in FOLFOX-resistance. Thus, downregulation of p21 can be an important mechanism for inducing TRAIL-mediated apoptosis in FOLFOX-resistant CRC cells. This development will impact patients across both genders and disparate groups afflicted by FOLFOX-resistant CRC. These studies may also be applied more broadly to other cancers and treatment paradigms where parallel PP2A decrease and AKT1/mTOR/4EBP1/p21-mediated acquired resistance is involved.

## Acknowledgements

This work was supported in part by the Department of Anatomy and Cell Biology and by Royalty Funds (#00126956), University of Florida, Gainesville, FL to S.N. A.K.S thanks the Department of Pharmacology, Penn State College of Medicine, and Penn State Cancer Institute (PSCI) for financial support. A.K.S. is also funded by NIH/NCI grant R21 CA234681 and CDMRP Lung Cancer Research Program (LCRP) Award Number W81XWH-19-1-0231. The authors thank Organic Synthesis Shared Resource of the Penn State Cancer Institute for synthesis of NSC49L. B.L. was supported in part by grants from the Florida Breast Cancer Foundation, the Ocala Royal Dames for Cancer Research, the Office of the Assistant Secretary of Defense for Health Affairs through the Breast Cancer Research Program under Award No. W81XWH-15-1-0199, and NIH/NCI grant CA252400.

## Author Contributions

S.N. developed the concept, performed experiments, and wrote the manuscript; A.K.S., synthesis of the NSC49L, was consistently involved during the development of the study; N.K. and M.A.R. performed experiments; I.M. and T.J.G., performed bioinformatic analysis; and B.K.L. and M.E.L., provided reagents and made intellectual contributions. All authors read the manuscript and provided their feedback.

## SUPPLEMENTARY FIGURES

**Figure S1.**
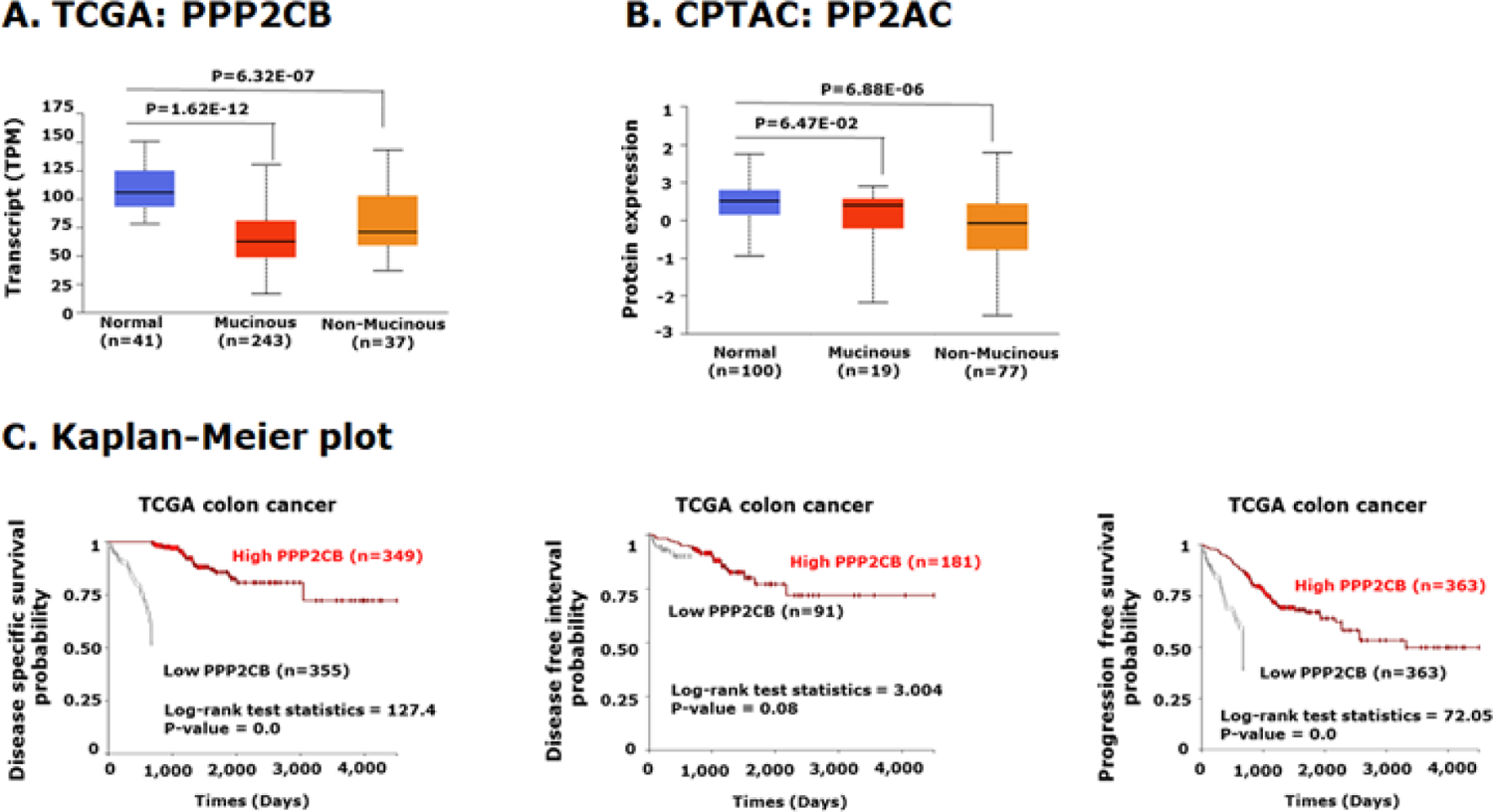
The mRNA and protein levels of PP2A in normal tissue and mucinous and non-mucinous CRC tumors. **A**, Box plot depicting a summary of TCGA data quantification of PPP2CB in normal tissue and CRC tumors (right), different stages (middle), and different nodal progression (left). **B,** Box plot showing the CPTAC data quantitation of PP2AC protein levels in normal tissues and CRC tumors (left), and different stages of progression (right). **C,** Kaplan-Meier survival analysis of TCGA data was plotted for the CRC cases that were high and low in PPP2CB expression. The p-value is shown on the top of each graph.

**Figure S2.**
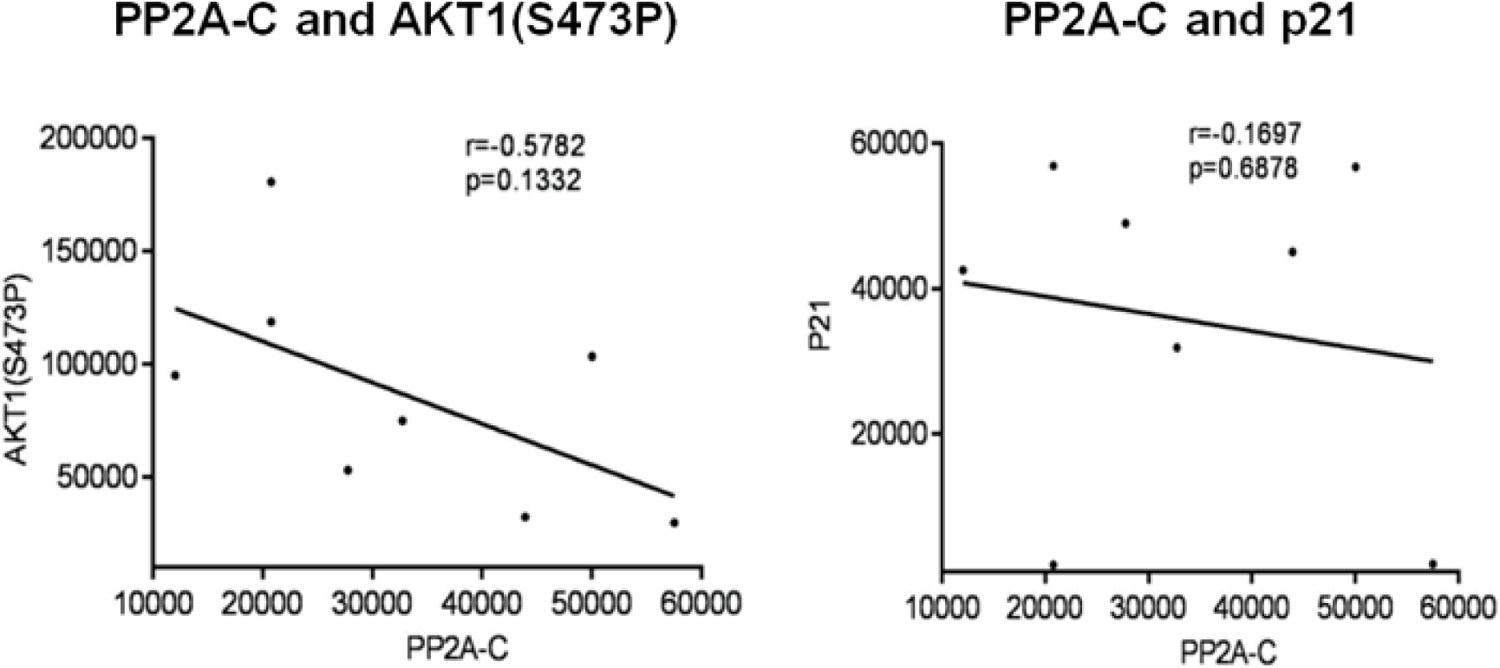
Correlation of PP2A expression level with AKT1(S473P) and p21 levels in CRC cell lines. To determine a correlation of PP2A with AKT1(S473P) and p21, we performed western bots of these proteins using the whole cell lysate of different CRC cell lines, including FOLFOX-resistant HCT116 and HT29 cells. A scatter plot of PP2A-C vs AKT1 (S473P) protein expression value from densitometry analysis of 8 cell lines generated with Graph Pad Prism 5 (La Jolla, CA, USA). Pearson correlation coefficients (r) and p-values are shown.

**Figure S3.**
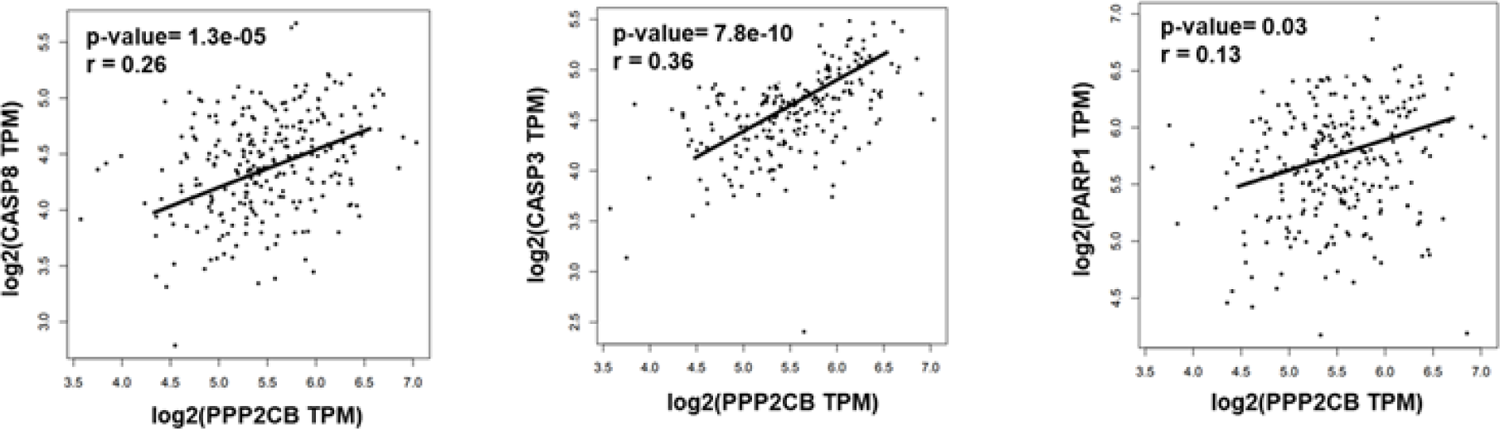
PPP2CB expression is positively correlated with the expression of caspase 8, caspase 3 and PARP1 levels in CRC tumors. We performed TCGA data analysis of PPP2CB and established a correlation with caspase 8, caspase 3 and PARP1 levels. Number of samples included in this analysis were 276.

**Figure S4.**
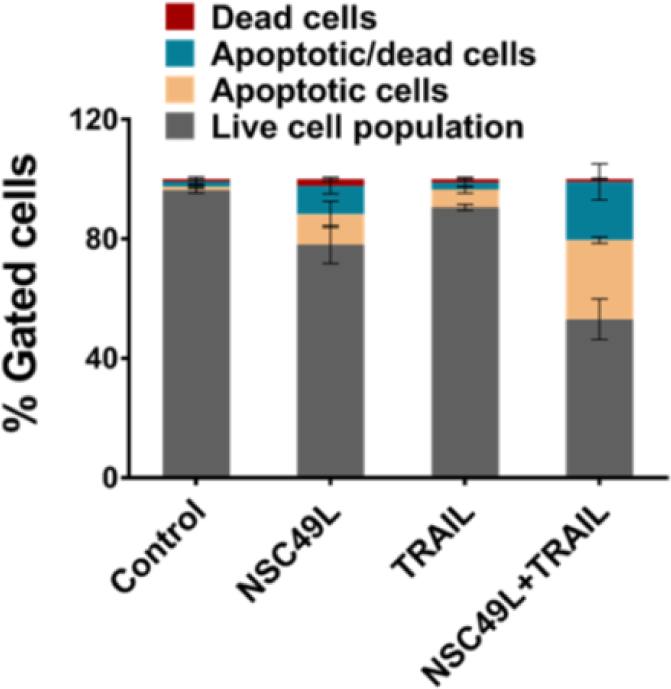
Measurement of apoptosis in FOLFOX-HCT116 cells treated with NSC49L and TRAIL. Cells were treated with NSC49L (20 µM) and TRAIL (1 nM) either alone or in combination for 24 h. The extent of apoptosis was determined by measuring the activation of caspase 3/7 using a flow cytometer. The experiment was repeated twice. Data are the mean ± SE of triplicate results.

**Figure S5.**
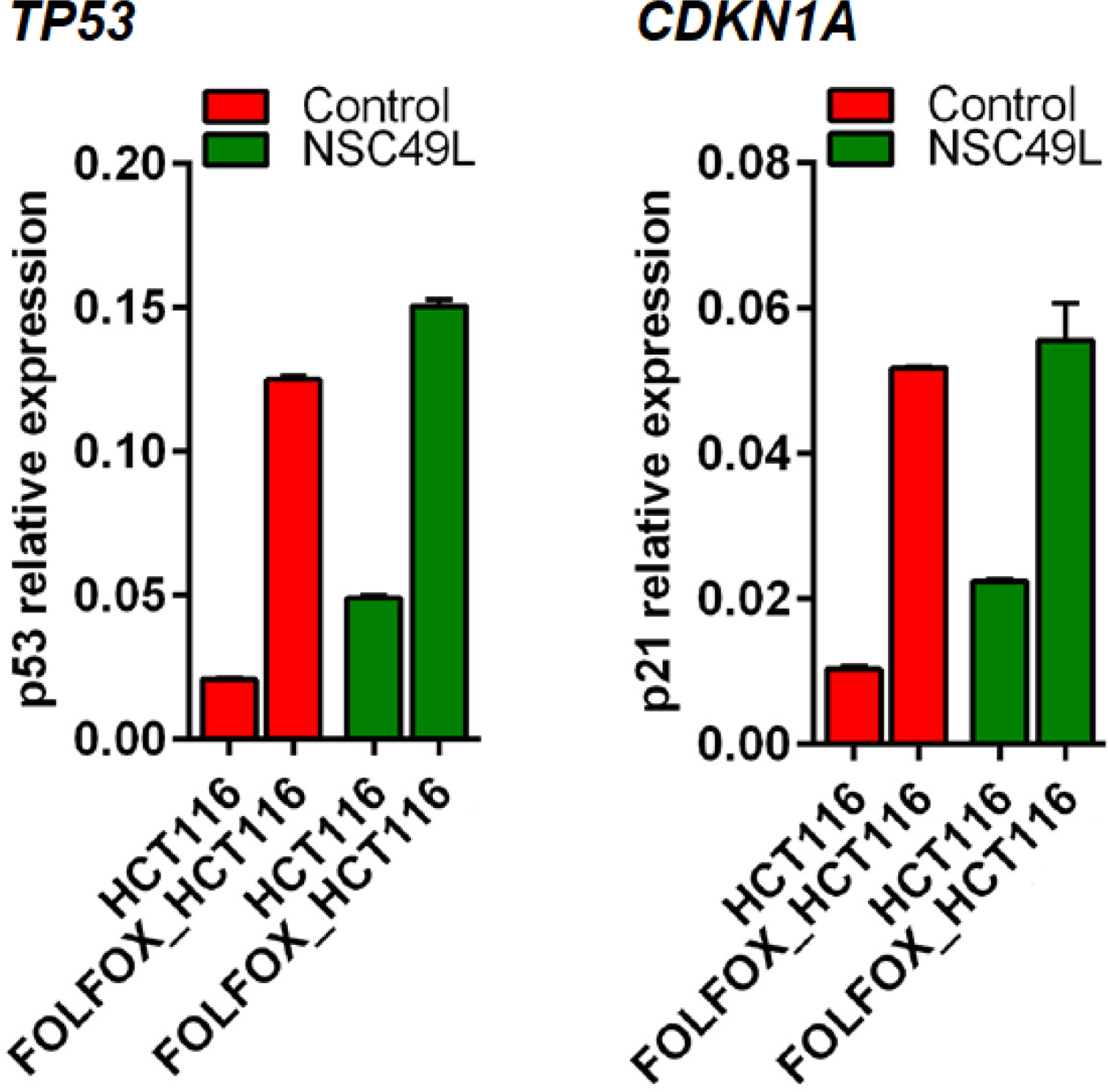
Effect of NSC49L and TRAIL treatments on the *TP53* and *CDKN1A* mRNA levels in HCT116 and FOLFOX-HCT116 cells. Cells were treated with NSC49L (20 µM) and TRAIL (1 nM) either alone or in combination for 16 h. Total RNA was isolated and used for qRT-PCR. Data are the mean ± SE of four estimations.

**Figure S6.**
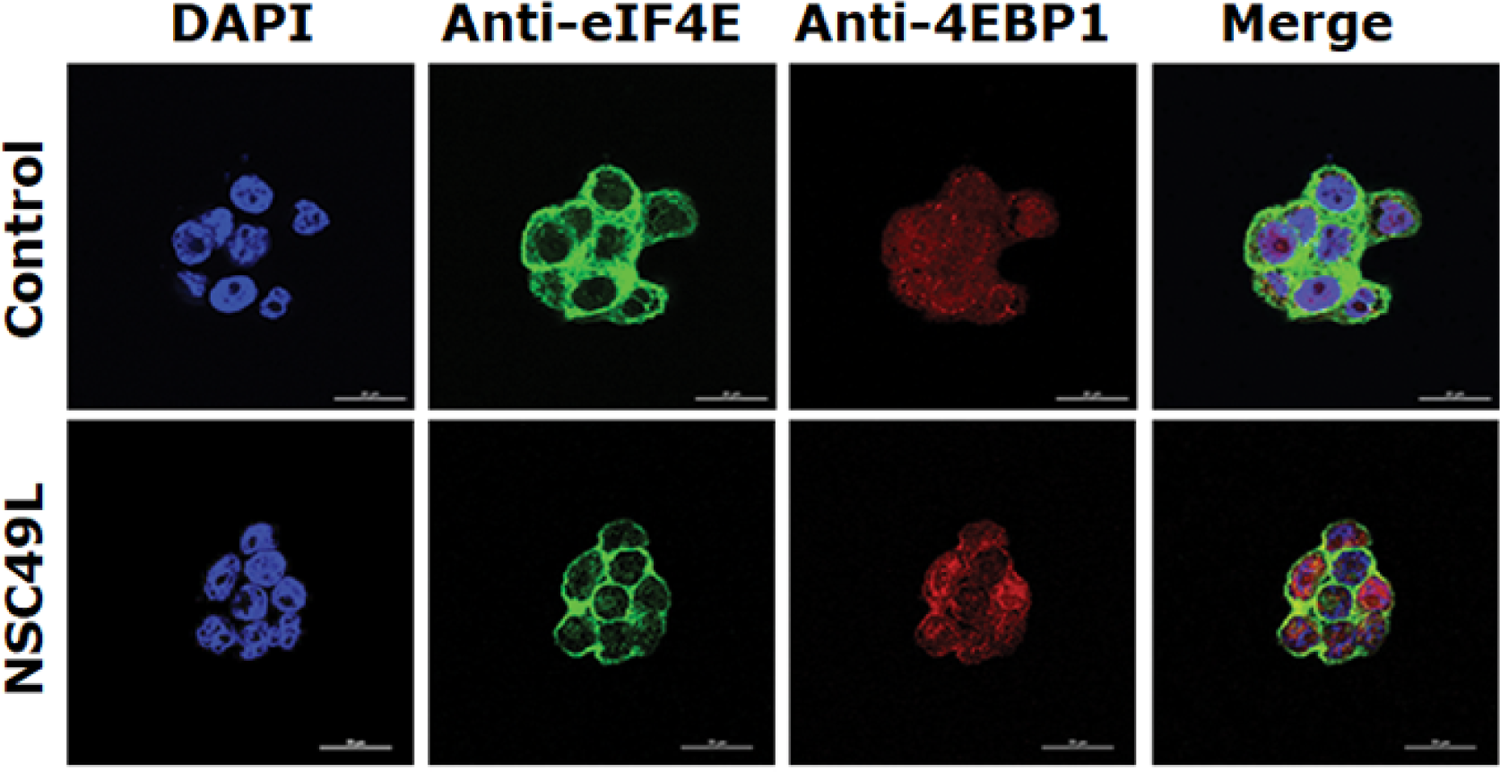
NSC49L and TRAIL treatments increase the colocalization of eIF4E and un-phosphorylated 4EBP1 in HT29 cells. HT29 cells were treated with NSC49L (20 µM) and TRAIL (1 nM) either alone or in combination for 24 h. After treatment, the colocalization of eIF4E and 4EBP1 was determined by IHC. Yellow color in merge column shows the colocalization of these two proteins. Images were captured at 60X magnification by Nikon A1RMPsi-STORM4.0 confocal microscope (Melville, NY).

**Figure S7.**
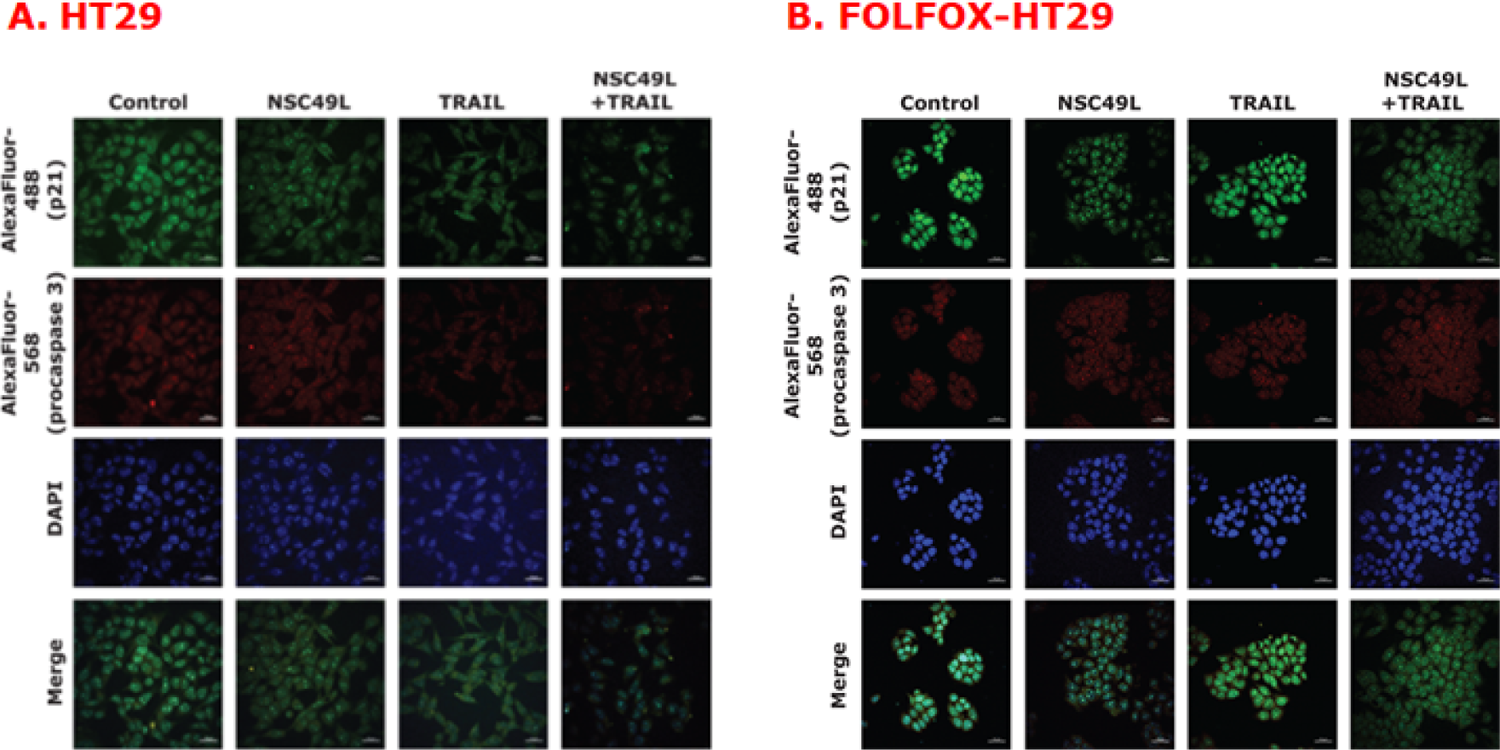
IHC analysis of p21 and procaspase 3 colocalization and levels in HT29 and FOLFOX-HT29 cells. Cells were grown on cover slips and treated with NSC49L (20 µM) and TRAIL (1 nM) either alone or in combination for 24 h and processed for IHC analysis. The secondary antibodies for p21 and procaspase 3 were AlexaFlour-488 (green) and AlexaFlour-568 (red), respectively. IHC images captured at 20X magnification by Leica DM IRBE confocal microscope (Hawthorn, NY).

## STAR*METHODS

- CRC cell lines
- Cell viability assay
- Apoptosis assay
- PP2A assay
- qRT-PCR
- RNAseq analysis
- Western blot analysis
- SUnSET assay
- Small molecule (NSC49L) synthesis
- Quantification and statistical analysis

### KEY RESOURCES TABLES

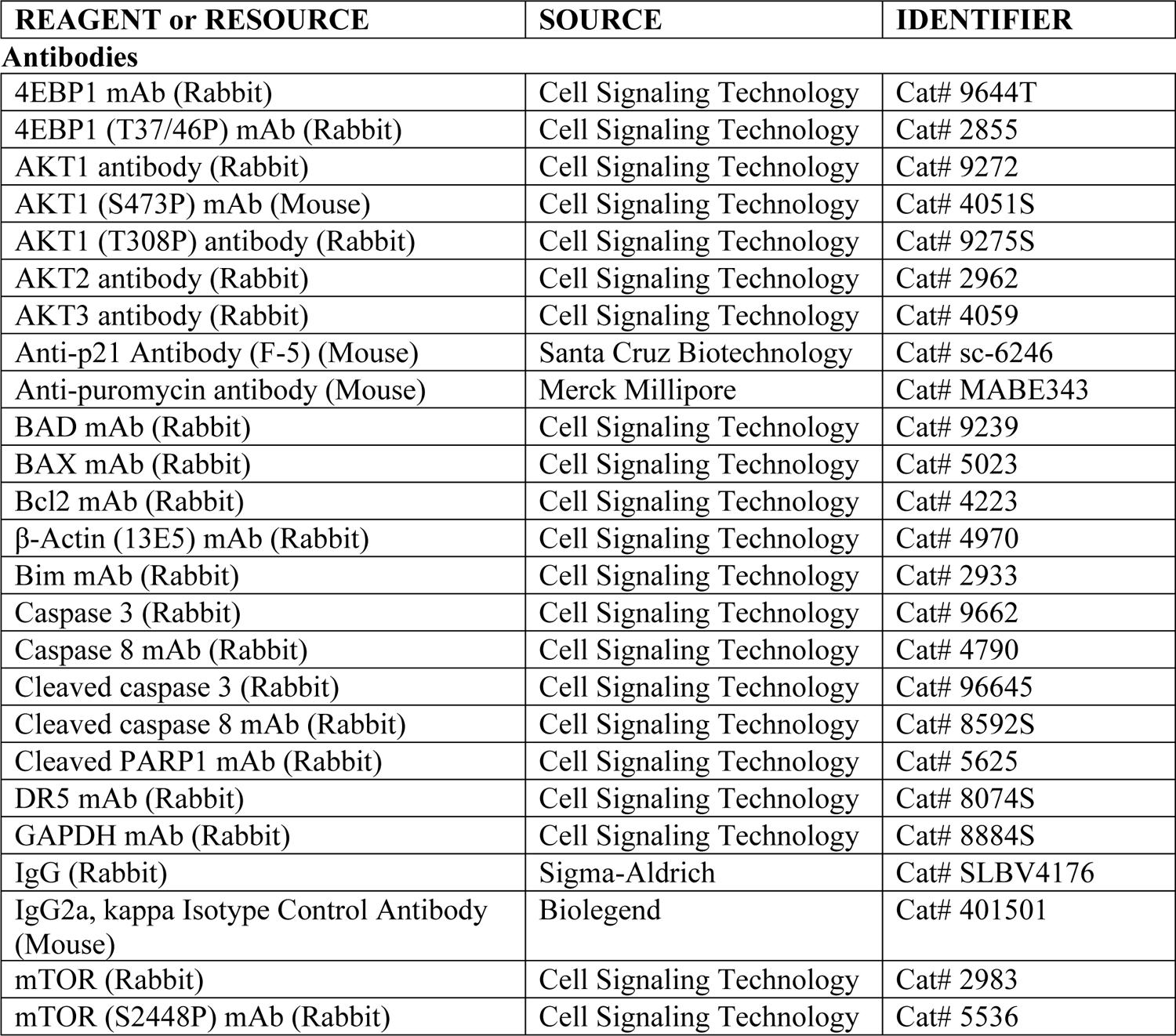
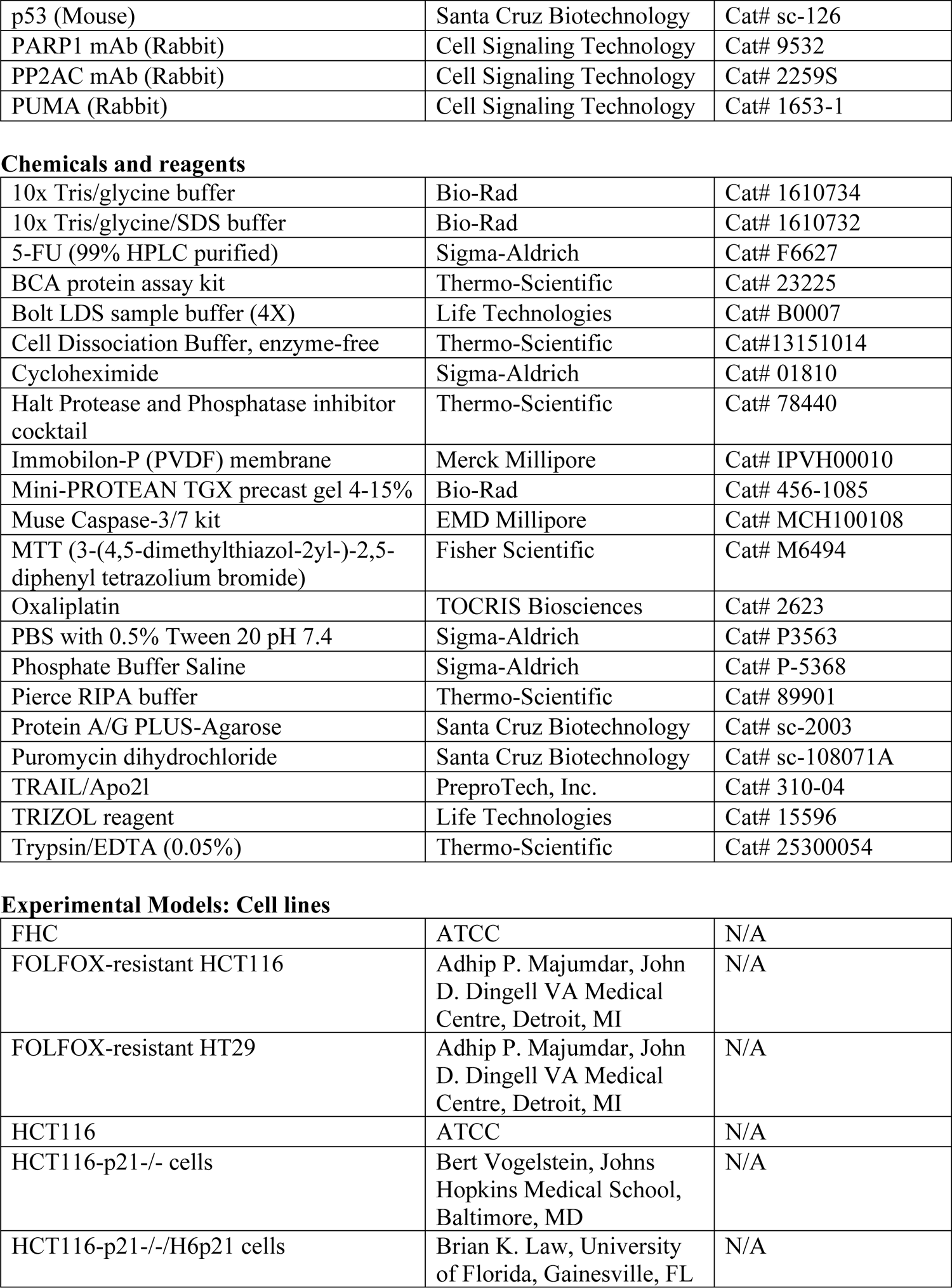
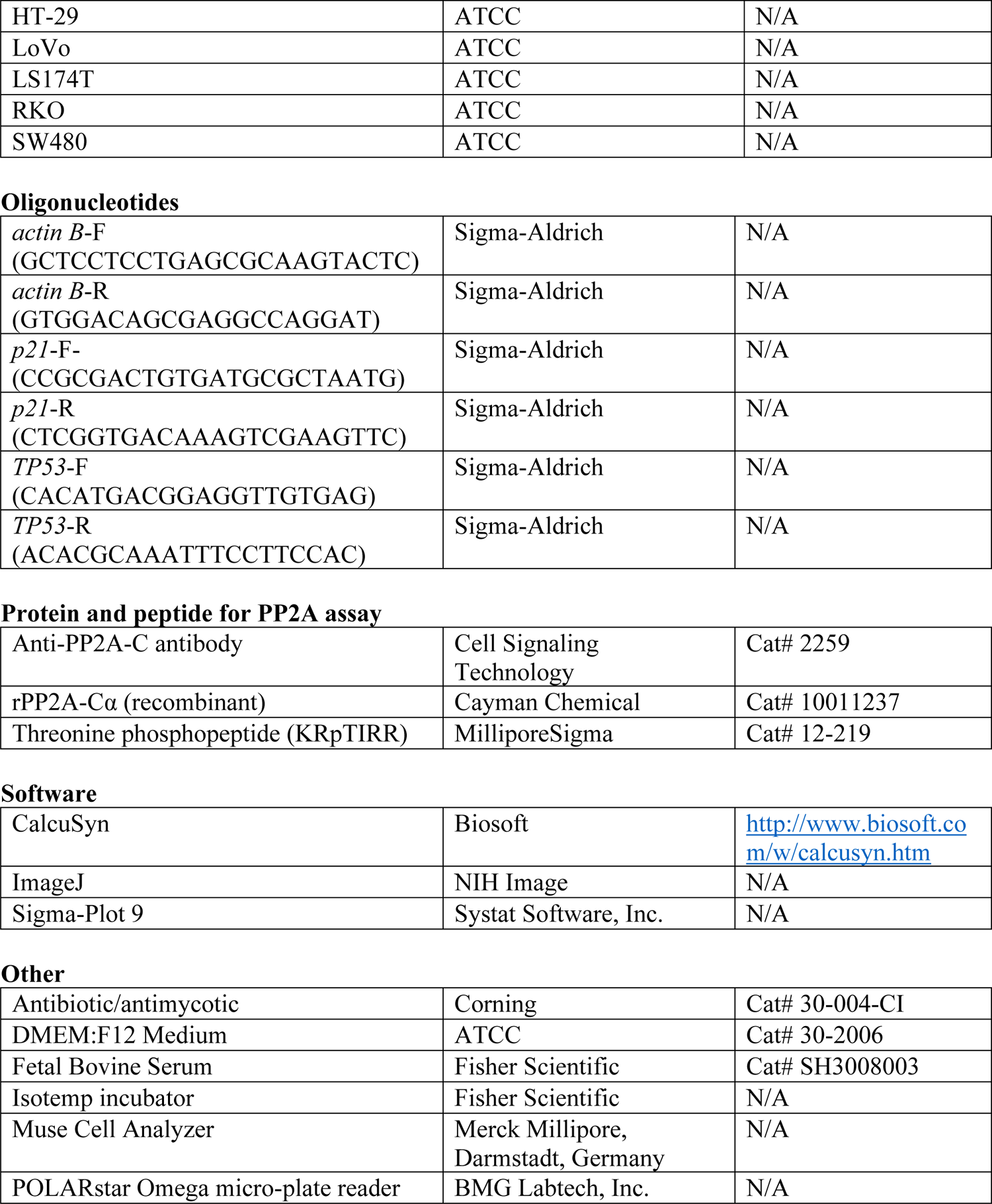

### RESOURCE AVAILABILITY

#### Lead Contact

Further information and requests for resources and reagents should be directed and will be fulfilled by the Lead Contact, Prof. Satya Narayan or Prof. Arun K. Sharma.

### Materials Availability

FOLFOX-resistant HCT116 and HT29 cell lines were obtained from Dr. Adhip P. Majumdar (John D. Dingell VA Medical Center, Detroit, MI). Other CRC and normal cell lines were obtained from ATCC. ATCC source information is provided in the Key Resources Table.

HCT116-p21^-/-^ cells were obtained from Dr. Bert Vogelstein (Johns Hopkins Medical School, Baltimore, MD). HCT116-p21^-/-^/H6p21 cells were created and characterized by Drs. Brian K. and Mary E. Law (University of Florida, Gainesville, FL) using previously described reagents and methods (Jahn et al., 2013; Law et al., 2006). HT29/tet-DR5 was constructed and characterized by Drs. Brian K. and Mary E. Law using the vectors and methods described previously (Wang et al., 2019). These cell lines can be requested from the above-mentioned investigators.

Synthesis of NSC49L was according to the previously published methods (Warmus, 1993). Compounds can be generated according to the described procedure or will be shared upon request by contacting Dr. Arun K. Sharma (Department of Pharmacology, Penn State University College of Medicine, Hershey, PA). Oxaliplatin and 5-FU were purchased from TOCRIS Biosciences and Sigma-Aldrich, respectively. TRAIL/Apo2L and PP2A-Cα were purchased from PreproTech, Inc. (Cranbury, NJ) and Cayman Chemical (Ann Arbor, MI), respectively.

## Data and Code Availability

TCGA data were for CRC’s are available at The Cancer Genome Atlas (TCGA) data repository (https://www.cancer.gov/about-nci/organization/ccg/research/structural-genomics/tcga/studied-cancers/colorectal). Gene expression data for FOLFOX-resistant and non-resistant patient are available at the NCBI Gene expression Omnibus (GEO) under GSE72970 accession (https://www.ncbi.nlm.nih.gov/geo/query/acc.cgi?acc=GSE72970).

## METHODS DETAILS

### CRC cell lines

Human colon cancer cell lines HCT-116, LoVo, SW480, LS174T, RKO, HT-29 and FHC were obtained from the American Type Culture Collection (ATCC, Rockville, MD). Cells were maintained as recommended by ATCC either in McCoy’s 5a or Dulbecco’s modified Eagle medium (DMEM; 4.5 g/L D-glucose) supplemented with 10% FBS and 1% antibiotic/antimycotic in tissue culture flasks in a humidified incubator at 37°C in an atmosphere of 95% air and 5% carbon dioxide. The medium was changed two times a week and cells were passaged using 0.05% trypsin/EDTA. The normal colonic epithelial cell line FHC was maintained in ATCC-formulated DMEM:F12 Medium Catalog #30-2006, containing 25 mM HEPES, 10 ng/ml cholera toxin, 0.005 mg/ml insulin, 0.005 mg/ml transferrin, 100 ng/ml hydrocortisone and fetal bovine serum 10% (v/v). The FOLFOX-resistant HCT-116 and HT29 cell lines were obtained from Dr. Adhip P. Majumdar (John D. Dingell VA Medical Center, Detroit, MI) (Yu et al., 2009), as described in (Narayan et al., 2017).

### Cell viability assay

The viability of cells was determined by MTT (3-(4,5-dimethylthiazol-2yl-)-2,5-diphenyl tetrazolium bromide) assay. In principle, the viable cell number is directly proportional to the purple formazan color of the reduced MTT dye, which can be quantitatively measured by spectrophotometry. Briefly, 1,500 cells were plated in quadruplets in 96-well flat-bottom tissue culture plates. After treatment with compounds for certain periods as described in respective figure legends, 10 µl of MTT reagent was added to each well and incubated at 37°C for 4 h to allow the formation of purple color crystals of formazan. In total, 100 µl of detergent solution was added to each well, and the reaction mixture was incubated in the dark for 2-4 h at room temperature. The developed color density was then measured spectrophotometrically at 570 nm using the POLARstar Omega micro-plate reader (BMG Labtech, Inc., Cary, NC).

### Apoptosis assay

The apoptosis assay using Muse Caspase-3/7 AAD assay kit was performed following manufacturer’s protocol. Briefly, cells were seeded in 6-well plate at a density of 1 x 10^5^ cells/well. Next day, cells were treated with 20 µM NSC49L and 1 nM TRAIL either alone or in combination for 24 h. Then, cells were harvested using enzyme-free Cell Dissociation Buffer.

The cells were resuspended in 1X Assay buffer BA. A 50 µL cell suspension and 5 µL Muse Caspase-3/7 reagent working solution was added in each tube. Samples were mixed thoroughly and gently by pipetting up and down, followed by incubation for 30 min at 37°C with 5% CO2. After incubation, 150 µL of Muse Caspase 7-AAD working solution was added to each tube and mixed thoroughly. Samples were incubated for 5 min in dark at room temperature, and then analysed in Muse Cell Analyzer. Events in each of the four quadrants were as follows. Lower-left: Viable cells, not undergoing detectable apoptosis [caspase-3/7 (-) and dead cell marker (-)]; Lower-right: cells in early stages of apoptosis [caspase-3/7 (+) and dead cell marker (-)]; Upper-right: cells in late stages of apoptosis or dead by apoptotic mechanism [caspase-3/7 (+) and dead cell marker (+)], and Upper-left: cells that have died via necrosis but not through apoptotic pathway [caspase-3/7 (-) and dead cell marker (+)].

### PP2A assay

We determined PP2A activity by using recombinant human PP2A-Cα protein and threonine phosphopeptide (KRpTIRR) substrate following the Malachite green protocol as previously described (Lek et al., 2017). The exact protocol we followed for the PP2A assay is given at: https://www.jove.com/pdf/55361/jove-protocol-55361-recombinant-synucleins-stimulate-protein-phosphatase-2a-catalytic. We used 0.0007 U/µl of human rPP2Ac, 0, 1, 10, 50, 100, 500 and 1,000 nM NSC49L, and 2 mM threonine phosphopeptide (KRpTIRR) substrate in this assay. PO4 standards (0, 150, 300, 600, 1,200, and 2,400 pmol) were run at the same time. Reaction was started by the addition of the Malachite green solution, incubated for 10 min at room temperature, and the change in color was read at 630 nm using POLARstar Omega micro-plate reader (BMG Labtech, Inc., Cary, NC).

We also determined PP2A activity in the immunocomplexes isolated from the FOLFOX-HT29 cells using anti-PP2A-C antibody as described (Lek et al., 2017). Cells were treated with NSC49L (0, 5, 10 and 20 µM) for 24 h. Whole lysates were prepared and immunocomplexes were isolated on Protein A Sepharose CL-4B beads. A 250 mg beads were diluted in 4 ml cold PBS containing 0.1% NaN3 (w/v), vortexed briefly, and incubated at room temperature in agitation (rotation) for 1 h. Then, the beads were spined down at 5,000 rpm for 1 min, washed 3-times with Co-IP buffer. Resuspend the beads in twice-volume of the Co-IP buffer. A 250 mg beads made 1.5 ml of the bed volume. Resuspend the pellet into 1.5 ml of Co-IP buffer with 1:1 ratio. Precleared the whole cell lysate by adding 100 µl of beads slurry (50 µl packed beads) per 100 µL of the whole cell lysate (containing 100 µg of protein) and incubated at 4°C for 10-30 min on a rocker or orbital shaker. Beads were removed by centrifugation at 14,000 x g at 4°C for 10 min. The supernatant was transferred to a fresh centrifuge tube and the pellet was discarded. To the 100 µg of precleared whole cell lysate, 2 µg Anti-PP2A, C subunit (52F8) rabbit monoclonal antibody was added and incubated at 4°C for 1-2 h or overnight on a rotator.

Immunocomplexes were captured by adding 50 µl Protein A sepharose CL-4B beads slurry (25 µl packed beads), and gently rocking on a rocker for 1 h or overnight at 4°C. The volume was brought to 500 µL with of ice-cold lysis buffer, and then centrifuged at 2,500 x g for 30 sec at 4°C. Carefully removed the supernatant and washed the beads 5-times with 500 µl of pNPP assay buffer. Finally, resuspended the beads into a pNPP buffer and determined the PP2A activity as described above.

### qRT-PCR

The mRNA levels of *CDKN1A* (*p21*) and *TP53* (*p53*) genes were determined by qRT-PCR. The total RNA was isolated from control and treated cells by using TRIZOL reagent. The nucleotide sequences used for the forward and reverse primers are described in the STAR*TABLE. The qRT-PCR was run on a real-time PCR machine (ABT 7500 fast) with standard cycle protocol. The relative mRNA abundance was determined using the differential cycle threshold (ΔCt) between the Ct values of cDNA from experiment and control cells.

### RNAseq analysis

For RNA-seq data analysis, we used the RNAseq data analysis pipeline reported previously. Briefly, Briefly, FASTQ files were aligned to Genome Reference Consortium Human Build 38 (GRCh38) using HISAT2; the transcripts assembling was performed using StringTie with RefSeq as transcripts ID; and the normalized counts (by FPKM) was called using Ballgown. The differential expression analysis was performed using R package limma; and the gene enrichment analysis was performed using R heatmap package.

### Western blot analysis

The levels of various proteins were determined by western blot analysis using 5-bromo-4-chloro-3-indolyl phosphate (BCIP)/nitro blue tetrazolium (NBT) substrate. Thirty micrograms of the whole cell lysate were separated by SDS-PAGE and transferred onto the PVDF membrane. After the transfer, the membrane was Incubated in the blocking solution (0.2 g/L NaN3, 50 g/L BSA and 2 ml/L Tween-20) for 60 min. Then, incubated with primary antibody (1:1,000, v/v in the blocking solution) at 4°C for overnight. Washed the membrane 3-times, each for 10 min, in a washing buffer ((Tris-base 6.05 g/L, CaCl2 (Dihydrate) 0.222 g/L, NaCl 4.75 g/L and Tween-20 500 µl/L, pH 8)). Incubated the membrane in a secondary antibody (1:5,000, v/v in the blocking solution) for 60 min. Washed the membrane 3-times, each for 10 min, in a washing buffer as described above. Layered the membrane with alkaline phosphatase substrate ((33 µl BCIP (5 mg/ml in 100% dimethylformamide) and 66 µl NBT (10 mg/ml in 70% dimethylformamide)) for 5-15 min. Once the desired signal appeared, the membrane washed 3-times with water. After membrane dries, the signal was scanned and recorded. In some cases, we used chemiluminescence method of western blotting as well.

### SUnSET assays

Following 24 h treatment with 20 µM of NSC49L, FOLFOX-HT29 cells were subjected to SUnSET assay (Schmidt et al., 2009), whereby cells were incubated with puromycin that was incorporated into the newly translated proteins. Briefly, after the onset of the treatment period, the cells were treated with vehicle control or with puromycin (10 µg/ml) or with puromycin and cycloheximide (CHX; 10 µg/ml) for 15 min (pulse). Further, the medium was removed, and the cells were washed thrice with fresh medium and incubated in fresh growth medium for an additional 2 h chase. The proteins were extracted, and 500 µg of cell lysate was used for immunoprecipitations (IP). The cell lysate was incubated with 5 µg of mouse control IgG2a (401501), K isotype, or 5 μg of mouse anti-puromycin antibody (MABE343) for 2 h at 4°C. Resuspended Protein A/G PLUS-Agarose (20 µl) was added to the cell lysate and mixed at 4°C overnight. Immunocomplexes were collected by centrifugation at 2,500 rpm for 5 min at 4°C. Pellets were washed 4-times with 1 ml PBS. After the final wash, the supernatant was discarded, and pellet was resuspended in 40 µl of 2X electrophoresis sample buffer. Proteins were then resolved by SDS-PAGE and analyzed by immunoblotting using an anti-p21 (F-5) antibody. The pre-immunoprecipitated lysate was used to assess the loading control, puromycin incorporation and p21 biosynthesis using β-actin antibody (1:1,000, v/v), anti-puromycin antibody (1:2,500, v/v) and p21 antibody (1:200, v/v), respectively.

### Small Molecule Synthesis

NSC49L was synthesized following the procedure of Warmus *et al*. (Narayan et al., 2019; Warmus, 1993). The final product was purified, crystallized, and characterized based on NMR and mass spectral data. The purity was determined to be at the level of ≥99%. Briefly, a mixture of 1,3,5,7-tetraazaadamantane (1.0 mmol) and 1 4-dichloro-2-butene (1.0 mmol) in CH2Cl2 (10 ml) was refluxed 6 h. After completion of the reaction as indicated by TLC, reaction mixture was cooled to room temperature, filtered, and the residue washed with excess methylene dichloride. The crude solid thus obtained was recrystallized in an EtOH/CH2Cl2 mixture and was dried under vacuum (1 mm) to afford NSC49L. Yield 92% (0.243 g); white solid; mp 199-201°C; ^1^H NMR (500 MHz, D2O) δ [ppm]: 6.31-6.27 (m,1H), 5.97-5.93 (m,1H), 5.08 (s, 6H), 4.71 (s, 3H), 4.54 (d, *J* = 11.0 Hz, 3H), 4.20 (d, *J* = 5.0 Hz, 2H), 3.57 (d, *J* = 6.0 Hz, 2H); ^13^C NMR (125 MHz, DMSO-*d6*) δ [ppm]: 140.6, 117.1, 78.1, 70.13, 57.8, 43.0; HRMS (ESI, M^+^) calcd. For C10H18ClN4 229.1219, found 229.1218.

## QUANTIFICATION AND STATISTICAL ANALYSIS

Statistical analyses for synergies between NSC49L and TRAIL treatments and p21 status were performed using CalcuSyn software (http://www.biosoft.com/w/calcusyn.htm). The combination index (CI) was calculated by applying Chou-Talalay method and was used for synergy quantification (Chou, 2010). Student’s *t*-test was used for comparisons in both *in vitro* and *in vivo* experiments. All experiments were repeated at least three times and results were expressed as mean ± SE. One-way analysis of variance (ANOVA) was calculated with Sigma-Plot 9. A one-tailed *t*-test was used to compare any significant differences between control and treated groups. The criterion for statistical significance was *p*<0.05. For western blotting data, band intensities were measured using ImageJ and normalized with GAPDH. Graph Pad Prism version 9.1.0 was also used for data visualization and p-value calculation.

